# Distinct Programs of Tissue Adaptation Shape Vitreous CD4^+^ and CD8^+^ T-cell States in Chronic Uveitis

**DOI:** 10.64898/2026.06.11.731765

**Authors:** Sandhya Rani Bhanja, Sayantan Ghosh, Jyotsana Negi, Arun Raina, Kaisar Alam, John V. Forrester, Pankaj Kumar, Soumyava Basu

**Affiliations:** Ocular immunology laboratory, Prof Brien Holden Eye Research Center, LV Prasad Eye Institute, Kallam Anji Reddy Campus, Hyderabad, India; Strand Lifesciences, Bangalore, India; Institute of Genomics and Integrative Biology, New Delhi, India; Department of Life Sciences, GITAM (deemed to be a university), Vishakhapatnam, India; Ocular Immunology Group, Section of Infection and Immunity, Institute of Medical Sciences, University of Aberdeen, Aberdeen, United Kingdom; Saroja A Rao Center for Uveitis, LV Prasad Eye Institute, Kallam Anji Reddy Campus, Hyderabad, India

## Abstract

Tissue-resident memory (T_RM_) T-cells are increasingly recognized as key mediators of chronic autoimmune inflammation, yet their organization and functional adaptation within the eye remain poorly understood. We investigated paired vitreous body biopsies and peripheral blood T cells from patients with chronic posterior segment uveitis using multiparameter flow cytometry, antigen-specific stimulation assays, single-cell RNA sequencing, and T-cell receptor sequencing to define the intraocular tissue-adaptive immune states. Vitreous T cells were phenotypically, transcriptionally, and clonally distinct from their circulating counterparts and enriched for canonical T_RM_ markers. However, tissue adaptation differed substantially between CD4^+^ and CD8^+^ lineages. Vitreous CD4^+^ T cells segregated into clonally expanded tissue-adaptive states characterized by greater CXCR6 expression, enhanced antigen-specific cytokine responses, and well-defined transcriptional profiles. In contrast, vitreous CD8^+^ T cells expressed higher levels of the retention-associated markers CD103 and CD49a yet maintained greater clonal and phenotypic continuity with peripheral blood T cells. Both vitreous CD4^+^ and CD8^+^ subsets exhibited a restrained effector profile associated with tissue-adaptive transcriptional programs. Our data reveal that CD4^+^ and CD8^+^ T cells in chronic uveitis assume distinct states in the vitreous microenvironment, such that the intraocular immune response relies on both localized tissue retention and active adaptation to the inflammatory niche.

## Main Text

Uveitis is a major cause of visual impairment and blindness, accounting for up to a fifth of all blindness worldwide (1). The disease course is often chronic or relapsing, leading to vision-threatening complications such as cataracts, glaucoma, macular edema, and others (2). Current therapeutic options, ranging from corticosteroids and conventional immunomodulatory therapy to targeted biologics, are effective at suppressing inflammation but have limited efficacy in preventing relapses or disease progression (3). Many patients receive prolonged immunosuppression without any substantial impact on long-term outcomes (4). Thus, there exists a critical need to better characterize intraocular immune responses in chronic or recurrent uveitis, beyond those in acute inflammation, which have been extensively studied in animal models (5,6).

Chronic inflammation is known to induce transcriptional, metabolic, and functional alterations in tissue-resident immune cells, thereby causing them to diverge phenotypically from their circulating counterparts (7-9). This phenomenon has been well described across multiple organs, including the skin, gut, lungs, and brain. These tissue-adapted states can persist long after the initial antigenic exposure, affecting both protective and pathogenic immunity. Tissue-resident memory (T_RM_) T cells remain the best characterized tissue-adapted immune cells (10,11). These have been classically described in CD8^+^ cells at barrier sites (skin, gut, lungs), with canonical markers such as CD69, CD103, CXCR6, and CD49a, which support their long-term retention in non-lymphoid tissues. The same markers have also been described in CD4^+^ T_RM_s, typically in non-epithelial zones, though less frequently than in CD8^+^ T_RM_s. Furthermore, in both CD4^+^ and CD8^+^ subsets, the expression of these markers is context-dependent and may vary across tissues and diseases. Tissue residency can be both under- and overestimated if defined solely by the expression of these markers (12,13). Advances in single-cell RNA and T-cell receptor sequencing (scRNA-seq and TCR-seq) have greatly expanded the range of transcriptional signatures available for identifying tissue-adapted cells across different organs (8,12,13). These insights highlight the need to combine phenotypic, functional, and molecular approaches to characterize tissue adaptation in chronic inflammation.

Despite these advances, the nature of tissue-adapted T-cell immunity within the eye remains poorly understood. The eye has historically been regarded as an immune-privileged organ, protected by the blood-ocular barriers and local immunosuppressive mechanisms (14). Yet the frequent chronicity of intraocular inflammation, both in eye-limited conditions and in association with systemic diseases, supports the likelihood of well-established resident immune populations in the inflamed eye (15). Prior work on ocular resident immune cells has largely focused on CD8^+^ cells in the corneal barrier tissue of animal models (16,17). In contrast, the vitreous body infiltrating cells in human uveitis, consisting of a heterogeneous mix of CD4^+^ and CD8^+^ cells, has yet to be studied for features of tissue adaptation. Unlike the rapidly circulating aqueous humor or cerebrospinal fluids (CSF), the vitreous is a low-turnover tissue compartment with resident hyalocytes and a collagen–hyaluronan extracellular matrix that prolongs the residence of infiltrating immune cells (18). This creates a unique microenvironment that could support tissue-adaptation programs for the cells trapped in the vitreous during chronic uveitis.

In this study, we sought to define the tissue-resident T cell subsets of the vitreous body in chronic posterior segment uveitis (PSU). We examined paired vitreous and peripheral blood samples from patients with PSU using multicolor flow cytometry and antigen stimulation assays, as well as scRNA-seq and TCR-seq. Our results show that vitreous-infiltrating CD4^+^ and CD8^+^ T cells are phenotypically, functionally, and transcriptionally distinct from their circulating counterparts. While CD4^+^ T cells segregated into discrete tissue-associated states within the vitreous, CD8^+^ T cells exhibited a more conserved transcriptional profile, maintaining substantial clonal and phenotypic overlap with their circulating counterparts despite acquiring tissue-retention features. Collectively, our study identifies distinct modes of tissue adaptation in vitreous CD4^+^ and CD8^+^ T cells and establishes the vitreous body as an immune-instructive niche that shapes local T-cell states during chronic uveitis.

## Results

### CD4^+^ and CD8^+^ T cells exhibit divergent modes of tissue adaptation within the vitreous

To determine whether T_RM_ T cells populate the vitreous compartment in patients with chronic uveitis, we performed unsupervised flow cytometry on paired vitreous and peripheral blood samples from patients with chronic PSU, as defined by the Standardization of Uveitis Nomenclature Working Group (19). While most (75.7%, n=54) patients were diagnosed with non-infectious uveitis (NIU), select phenotypes of tuberculosis (TB)-associated uveitis (e.g., intermediate uveitis or panuveitis) were also included (24.3%, n=18), as they did not reveal any overt granuloma in the eye and were phenotypically similar to NIU. Furthermore, our earlier studies have demonstrated autoreactive T-cell responses in the vitreous in both infectious and non-infectious uveitis, suggesting that despite differences in disease etiology, intraocular inflammation is sustained by common adaptive immune mechanisms (20-22). The samples were collected during diagnostic or therapeutic vitrectomy surgery and processed as described previously (23). Although the normal vitreous is generally devoid of immune cells, posterior segment inflammation involving the retina and choroid is typically associated with infiltration of immune cells into the vitreous, clinically recognizable as vitritis. Thus, the vitreous compartment, like other fluid-containing spaces, such as the CSF space or the joint space (24,25), serves as an indicator of tissue inflammation in the eye. We have previously reported the broad surface and functional phenotypes of vitreous-infiltrating immune cells in various posterior segment uveitides, with T cells accounting for >80% of the infiltrate (20,21). However, the tissue residency features of these cells have not been characterized.

The total immune cells were gated on singlet and live CD3^+^ population. The markers were selected for subset (CD4, CD8), memory T-cell (CD62L, CD45RA) and tissue residency (CD69, CD103) identification. CD4^+^ and CD8^+^ cells were analyzed separately using unsupervised dimensionality reduction and clustering. We identified 12 distinct clusters among CD4^+^ cells across the blood and vitreous compartments (Figure 1A - C). Five of these clusters were significantly enriched in the vitreous, and five in the blood. No significant difference was observed between blood and vitreous in two clusters. All vitreous-enriched clusters lacked expression of naïve and central memory markers CD45RA and CD62L and variably expressed tissue-residency markers CD69 and CD103, pointing to their T_RM_ background (6) (Figure 1D). Among these, cluster 6, the largest vitreous-enriched cluster, expressed both CD69 and CD103. Cluster 11, the most abundant blood cluster, retained CD62L and lacked CD45RA, consistent with a central memory T cell (T_CM_) cluster (26). Cluster 12, which was comparably represented in both vitreous and blood, lacked expression of CD45RA, CD62L, as well as CD69, and CD103 (Figure 1 E), suggestive of a recirculating effector memory (T_EM_)-like population rather than a tissue-resident state. Clusters 3 and 9, also notably absent from vitreous, expressed both CD45RA and CD62L, suggesting a naïve T-cell population.

**FIGURE 1:**
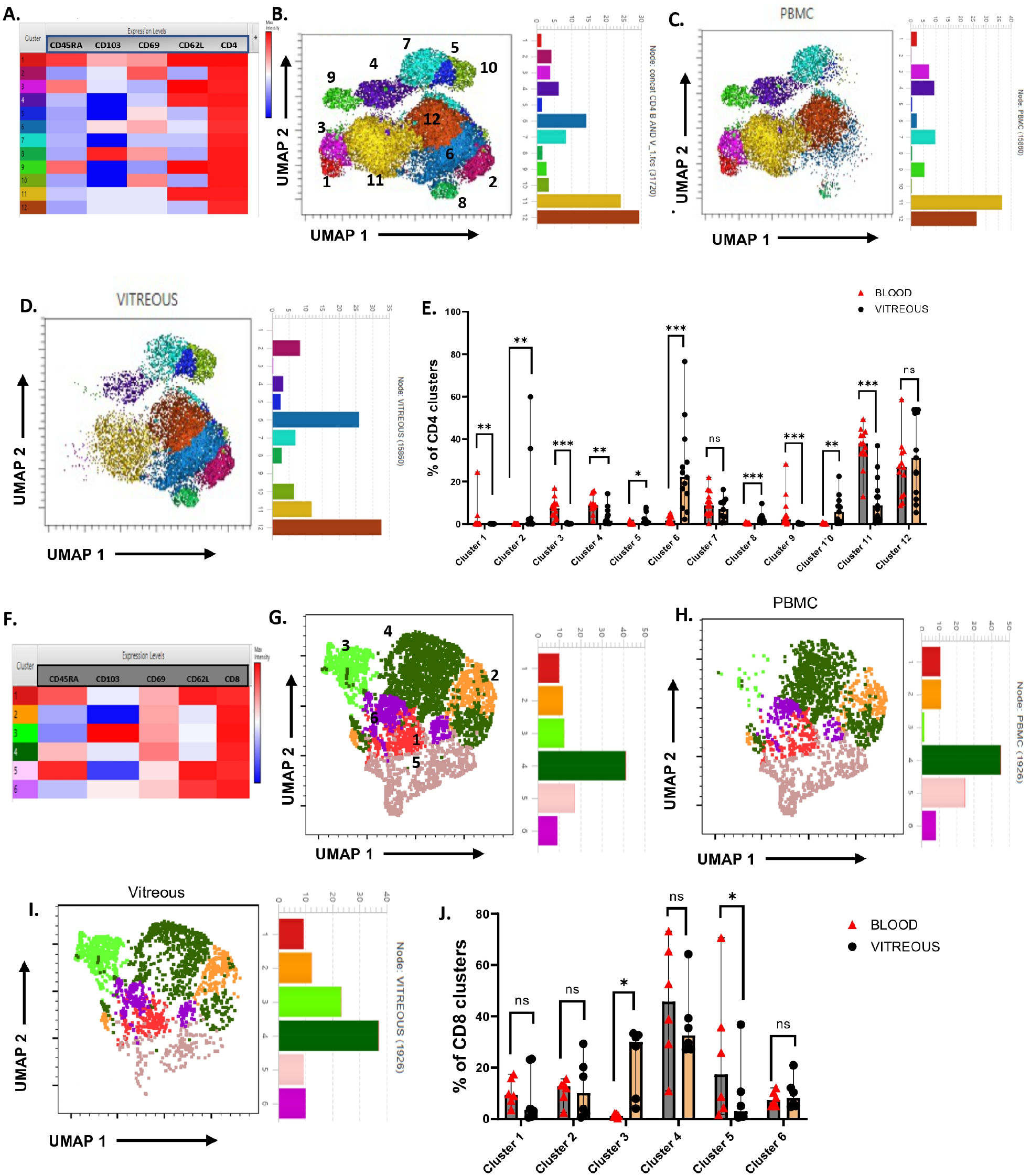
Unsupervised flow cytometry analysis identifies a phenotypically unique T cell population in vitreous compartment. **A**. Heatmap showing relative expression of the indicated markers in each of the CD4+ clusters. **B**. Combined UMAP analysis and clustering of 31720 CD3CD4+ T-cells from blood and vitreous of uveitis patients based on the expression of CD4, CD62L, CD45RA, CD103 and CD69. Bar plot on the right side of each UMAP plot indicates the relative size of clusters.**C**. unsupervised clustering of only PBMC. **D**. unsupervised clustering of only vitreous. **E**. Frequencies of CD4+ clusters in blood versus vitreous (n=13). Statistical analysis was performed using multiple t test (Wilcoxon sign) with Benjamini-Hochberg correction. **F**. Heatmap showing relative expression of the indicated markers in each of the CD4+ clusters. **G**. Combined UMAP analysis and clustering of 3852 CD8+ T-cells from blood and vitreous of uveitis patients based on the expression of CD4, CD62L, CD45RA, CD103 and CD69. Bar plot on the right side of each UMAP plot indicates the relative size of clusters. **H**. unsupervised clustering of only PBMC. **I**. unsupervised clustering of only vitreous. **J**. Frequencies of clusters in blood versus vitreous in CD8 clusters (n=16). Statistical analysis was performed using paired multiple t test (Wilcoxon sign) with Benjamini-Hochberg correction. Individual *P* values are noted on respective graphs or summarized as: \**P* < 0.05, \*\**P* < 0.01, \*\*\**P* < 0.001, \*\*\*\**P* < 0.0001. Bars show median ± interquartile ring. High dimensional data were created using FlowJo v10 software.

In contrast to CD4^+^ T cells, unsupervised analysis of CD8^+^ T-cells revealed greater phenotypic overlap between the blood and vitreous compartments (Figure 1 F - I). Six CD8^+^ clusters were identified, only two of which showed significant compartmental enrichment (Figure 1 J). Cluster 3, which was relatively enriched in the vitreous, was characterized by loss of CD45RA and CD62L while expressing both CD69 and CD103. Conversely, cluster 5 was a blood-enriched cluster that expressed CD45RA and CD62L, along with low levels of CD69, consistent with a recently activated naïve or transitional phenotype. Together, our findings suggest that while the vitreous CD4^+^ cells adopt differential T-cell subsets in chronic uveitis, CD8^+^ cells generally show phenotypic continuity between blood and vitreous, providing a framework for subsequent functional and transcriptional analyses.

### Canonical T_RM_ markers are heterogeneously expressed by vitreous CD4^+^ and CD8^+^ T-cells

Next, we sought to characterize the distribution of canonical T_RM_ markers in vitreous samples from patients with chronic uveitis. Our earlier studies have revealed that the vitreous-infiltrating cells in chronic PSU primarily comprise memory T cells (20, 21). Among these, T_EM_ cells were the major memory subtype, much like other sites of tissue inflammation (Figure 2A) (27). We argued that vitreous memory T-cells in chronic uveitis will acquire tissue-residency characteristics as they remain confined to the vitreous compartment for prolonged periods. We selected CD69 and CD103 as the primary tissue residency markers for both CD4 and CD8 cells, in line with current norms (7,10,11,28). While CD103 has classically been associated with epithelium-localized CD8^+^ T_RM_ cells, it has also been reported in certain stromal tissues, such as the lamina propria of the small intestine (29), and in CD4^+^ cells in human lungs (30). Phenotypic analysis of vitreous T-cells (n = 31) revealed distinct patterns of CD69 and CD103 expression across CD4^+^ and CD8^+^ T cell compartments that were gated negatively for CD45RA and CD62L expression (Figure 2B). Among CD4^+^ T-cells, the CD69^+^CD103^−^ memory subset represented the predominant population compared with CD69^−^CD103^−^, CD69^+^CD103^+^, and CD69^−^CD103^+^ subsets (P < 0.0001) (Figure 2C). In contrast, CD8^+^ T-cells displayed significantly higher frequencies of both CD69^+^CD103^−^ and CD69^+^CD103^+^ subsets compared with CD69^−^CD103^−^ cells (P < 0.0001), with CD69^+^CD103^+^ cells comprising a substantially larger fraction of the CD8^+^ compartment than observed among CD4^+^ T-cells (Figure 2D). While CD69 expression alone could also denote early activation in CD4^+^ T_EM_ cells (31), we found that the CD69^+^CD103^−^ cells expressed lower levels of another activation marker, CD38 (32), than the CD69^+^CD103^+^ cells (Figure 2 E - G). This suggests that CD69 expression alone, could represent a T_RM_ phenotype within the vitreous microenvironment rather than merely a marker of early activation (6,8,25). Finally, sample-wise paired analysis confirmed significantly greater CD103 expression in CD8^+^ compared with corresponding CD4^+^ T-cells (P < 0.0001) (Supplemental Figure 1A), indicating preferential enrichment of CD103^+^ T_RM_-like cells within the vitreous CD8^+^ T cell pool. Taken together, we inferred that both CD4^+^ and CD8^+^ cells express canonical T_RM_ markers in the vitreous, though CD103 may be preferentially expressed on CD8^+^ cells.

**FIGURE 2:**
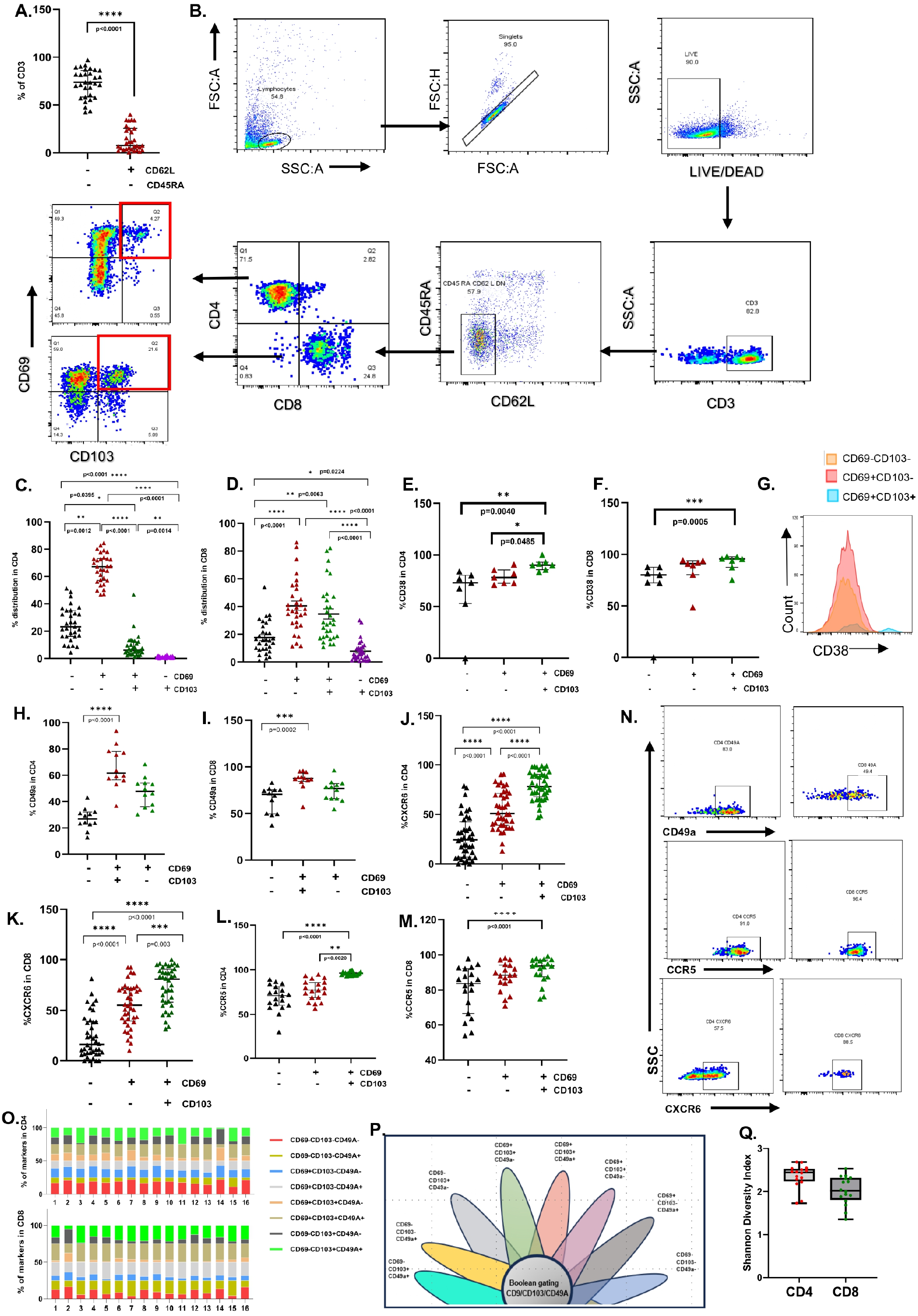
Heterogenous pattern of tissue residency markers in vitreous CD4 and CD8 T-cells. **A**. T effector memory cells (CD45RA-CD62L-) are high in vitreous compartment (n=31). **B**. Gating strategy applied to analyse tissue residency markers. **C-D**. T_RM_ distribution in CD4+ and CD8+ T-cells(n=31). Kruskal-Wallis with Dunn’s test as post hoc analysis is used. **E-G**. CD38 in CD4 and CD8 (n=7). Friedman test for paired data with Dunn’s test as post hoc analysis is used for comparison. **H-I**. CD49a expression in CD4 and CD8 (n=12). Representative flow cytometric analysis of CD49a expression. Friedman test for paired data with Dunn’s test as post hoc analysis is used for comparison. **J-K**. CXCR6 expression in CD4 and CD8 (n=40). Representative flow cytometric analysis of CXCR6 expression. Friedman test for paired data with Dunn’s test as post hoc analysis is used for comparison. **L-N**. CCR5 expression in CD4 and CD8 (n=19). Representative flow cytometric analysis of CCR5 expression. **O**. Stacked graph showing diversity of tissue residency marker expression in vitreous infiltrating T-cells in 16 patients showing intra heterogeneity. **P-Q**. Boolean gating strategy used to get different combination of tissue residency markers. Heterogeneity is measured by calculating Shannon index diversity for both CD4 and CD8 independently (n=16). Each point represents an individual patient sample with box plot showing median with interquartile range. Individual *P* values are noted on respective graphs or summarized as: \**P* < 0.05, \*\**P* < 0.01, \*\*\**P* < 0.001, \*\*\*\**P* < 0.0001. Bars show median ± interquartile range.

We next examined the distribution of the tissue-retention marker CD49a across CD69/CD103-defined T_RM_ subsets (12, 33). Although infrequently described in the context of CD4^+^ T_RM_ cells (34), CD49a has been identified as a marker of cytotoxicity and IFNγ production in CD8^+^ T_RM_ cells (33). We found that in CD4^+^ T-cells, CD49a expression was significantly enriched in CD69^+^CD103^+^ cells compared with CD69^−^CD103^−^ cells (p< 0.0001), with intermediate expression in CD69^+^CD103^−^ subsets (Figure 2H). A similar pattern was observed in CD8^+^ T-cells, where CD69^+^CD103^+^ cells exhibited significantly higher frequencies of CD49a expression (p=0.0002) (Figure 2I). However, sample-wise paired analysis revealed greater CD49a expression in CD8^+^CD69^+^CD103^+^ cells compared with corresponding CD4^+^ T-cells (Supplemental Figure 1B). Next, we examined the expression of chemokine receptors CXCR6 and CCR5 to support the tissue adaptation of CD69^+^ subsets. CXCR6 expression was significantly enriched within CD69^+^ populations in both CD8^+^ and CD4^+^ compartments (Figure 2 J - K), with the highest frequencies observed in CD69^+^CD103^+^ cells (p< 0.0001). Similarly, CCR5 expression was markedly elevated within CD69^+^CD103^+^ subsets in both CD4^+^ and CD8^+^ T-cells compared with CD69^−^CD103^−^ populations (P < 0.0001), with the highest expression consistently observed in CD69^+^CD103^+^ cells (Figure 2 L - N). However, unlike CD103 and CD49a, on sample-wise paired analysis, greater CXCR6 expression was noted in the CD4^+^CD69^+^CD103^+^ cells compared with corresponding CD8^+^ T-cells (Supplemental Figure 1C). No such difference was noted for CCR5 expression (Supplemental Figure 1D).

Together, these findings demonstrate that CD8^+^ T-cells preferentially expressed retention-associated T_RM_ markers (CD103 and CD49a), whereas CD4^+^ T-cells demonstrated stronger enrichment of inflammatory tissue homing programs (CXCR6).

Finally, we examined whether T_RM_ markers are consistently expressed across all patients with chronic uveitis. To do this, we used a Boolean gating strategy to analyze combinations of CD69, CD103, and CD49a expression. We included CD49a because it is predominantly retained in tissues and has been shown to perform better than other T_RM_ markers, such as CXCR6 in parabiosis experiments (12). This yielded eight distinct phenotypic subsets (2^3^=8) within the CD4^+^ and CD8^+^ populations in sixteen patients, respectively. We then assessed phenotypic heterogeneity in these populations by calculating the Shannon Diversity Index based on the frequencies of the eight Boolean-defined subsets. Both CD4^+^ and CD8^+^ T cells exhibited a range of these phenotypes across all 16 patients. Stacked bar plots revealed variability in the distribution of these populations across samples, indicating differences in tissue-adapted and other T-cell phenotypes among individuals (Figure 2O-P). CD4^+^ T cells showed higher diversity, with mean Shannon indices of around 2.4, compared to 2.0 for CD8^+^ T cells, suggesting moderate to high phenotypic heterogeneity (Figure 2 Q). This also indicates that CD4^+^ cells were more evenly distributed across the eight Boolean subsets and occupied a broader spectrum of tissue-adaptive states, whereas CD8^+^ cells clustered into a few dominant phenotypes, with some cells not expressing T_RM_ markers at all. These findings suggest that while canonical T_RM_ markers identify tissue-adapted populations in the vitreous, they do not reflect the full range of immune adaptation present in chronic ocular inflammation.

### Antigen-specific cytokine responses localize to CD69^+^CD103^+^ CD4 T-cells, while CD8 subsets retain polyclonal effector capacity

T_RM_ cells have distinct functional specializations at sites of inflammation, though these have largely been described in mouse models (10,11). While CD4^+^ T_RM_ cells exert helper and immunoregulatory functions through rapid cytokine production (30), CD8^+^ T_RM_s are more specialized for direct cytotoxicity while also producing proinflammatory cytokines (33,34). We first evaluated the intrinsic effector capacity independent of antigen specificity by stimulating vitreous T-cells with anti-CD3/CD28 beads. Here, CD4^+^ T-cells produced significantly higher TNFα and IL-17A in the CD69^+^CD103^+^ subsets compared to the CD69^−^CD103^−^ and CD69^+^CD103^−^ subsets (p<0.001–0.0001) (Figure 3 A - C). IFN-γ production was also enriched within CD69^+^CD103^+^ CD4^+^ T-cells relative to CD69^−^ subsets (p<0.05) (Figure 3 D) while no obvious difference was noted for granzyme B (Gzmb) (Supplemental Figure 2A). In the CD8^+^ compartment, polyclonal stimulation induced IFN-γ and TNF-α production across multiple subsets, with significant enrichment observed in CD69^+^CD103^+^ cells compared to the non-T_RM_ population (p<0.01–0.001) (Figure 3 F - H). No difference was noted between groups for IL-17a (Figure 3 E) or Gzmb (Supplemental Figure 2B-J), though a trend towards increased production of Gzmb by CD69^+^CD103^+^ cells was noted. In both CD4^+^ and CD8^+^ populations, the CD69^+^CD103^+^ subset also had higher polyfunctional response when evaluated for all three cytokines (TNFα, IFNγ and IL-17a) or the TNFα, IFNγ combination (Figure 3 I – J). Non-stimulated samples also had elevated cytokine levels, likely due to their origin from the inflamed site. These data indicate that T_RM_-like T-cells in the vitreous retain substantial intrinsic effector capacity upon TCR engagement.

**FIGURE 3:**
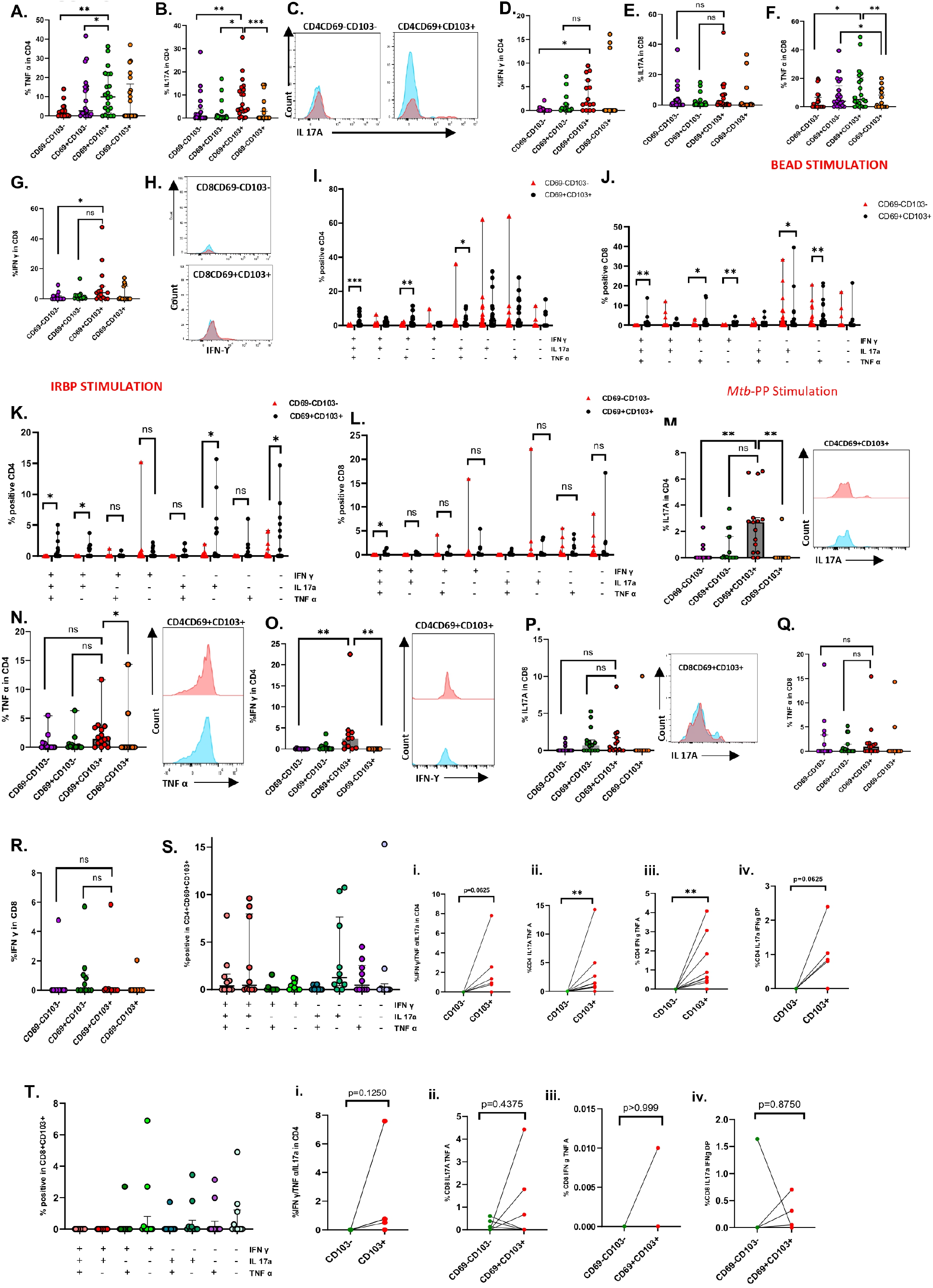
Antigen-specific stimulation shows monofunctional and polyfunctional cytokine profile in inflamed uveitis vitreous T-cells. **A**-**D**. Monofunctional response of TNFα (n=21), IL-17A (n=21), IFN-γ (n=15) and Gzmb (n=15) in CD4 T-cells upon CD3/CD28 TCR mediated stimulation. Friedman test for paired data with Tukey’s test as post hoc analysis. **E-H**. Monofunctional response of TNFα (n=21), IL-17A (n=21), IFN-γ (n=15) and Gzmb (n=15) in CD8 T-cells upon CD3/CD28 TCR mediated stimulation. Friedman test for paired data with Dunn’s test as post hoc analysis. **I - J**. Polyfunctional response comparison of TNFα, IL-17A, IFN-γ in CD69-CD103- and CD4CD69+CD103+ subsets (n=20). **K-L**. Polyfunctional response comparison of TNFα, IL-17A, IFN-γ in CD69-CD103-and CD4CD69+CD103+ subsets upon IRBP stimulation (n=8). **M-O**. Monofunctional response of TNFα (n=15), IL-17A (n=15) and IFN-γ (n=11) in CD4 T-cells upon *Mtb*-PP stimulation. Friedman test for paired data with Dunn’s test as post hoc analysis is used for comparison. **P-R**. Monofunctional response of TNFα (n=15), IL-17A (n=15) and IFN-γ (n=11) in CD8 T-cells upon *Mtb*-PP stimulation. Friedman test for paired data with Dunn’s test as post hoc analysis is used for comparison. **S**. Quantification of IFN-γ, TNF-α, and IL17a co-expression in CD4+CD69+CD103+ subset (n=10). **Si-iv**. Polyfunctional response comparison of TNFα/IL-17A/IFN-γ, IL17a/TNFα, IFN-γ/TNFα and IL17a/IFN-γ in CD4+CD69-CD103- and CD4+CD69+CD103+ subsets. **T**. Quantification of IFN-γ, TNF-α, and IL17a co-expression in CD8+CD69+CD103+ subset (n=10). **T i-iv**. Polyfunctional response comparison of TNFα/IL-17A/IFN-γ, IL17a/TNFα, IFN-γ/TNFα and IL17a/IFN-γ in CD8CD69-CD103- and CD8CD69+CD103+ subsets. For polyfunctional comparison multiple t test (Wilcoxon sign) with Benjamini-Hochberg correction is done. Individual *P* values are noted on respective graphs or summarized as: \**P* < 0.05, \*\**P* < 0.01, \*\*\**P* < 0.001, \*\*\*\**P* < 0.0001. Bars show median ± interquartile ring. In representative histogram images blue peak is represents non stimulated while pink represents the stimulated sample.

Antigen-specific functional responses have not generally been characterized in human T_RM_ studies. We have previously shown both monofunctional and polyfunctional responses from vitreous T cells of uveitis patients to the retinal autoantigen, interphotoreceptor retinoid-binding protein (IRBP, 20), and to *the Mycobacterium tuberculosis* peptide pool (*Mtb*-PP) consisting of ESAT-6(Early Secretory Antigenic Target-6) and CFP-10(culture filtrate protein-10) peptides, in patients with ocular TB (20-22). However, the contributions of T_RM_-specific responses were not examined. Here, stimulation of vitreous T-cells with IRBP revealed significantly greater polyfunctional responses in the CD69^+^CD103^+^ subset, which were more pronounced in the CD4^+^ compartment (Figure 3K-L). Monofunctional IL-17a, IFNγ, TNFα, and Gzmb responses were similarly increased in the CD4^+^CD69^+^CD103^+^ cells (Supplemental Figure 2 K-N), though not in the CD8^+^ subset (Supplemental Figure 2 O - R). A similar pattern was observed following Mtb-PP stimulation in ocular TB (as per the Collaborative Ocular Tuberculosis Study diagnostic criteria) (35). CD4^+^CD69^+^CD103^+^ cells demonstrated significantly enhanced IFNγ, TNFα, and IL-17A monofunctional responses (Figure 3M–O) lineage (Supplemental Figure 3 C-J), along with increased IL-17A–TNFα and IFNγ–TNFα polyfunctionality, whereas no significant enrichment was observed within CD8+ subsets (Figure 3P–T). No significant differences in Gzmb responses were detected in either lineage (Supplemental Figure 3 A-B). Together, these findings suggest that while both CD4^+^ and CD8^+^ T-cells with T_RM_ phenotypes are primed for effector responses, antigen-specific inflammatory specialization is concentrated among tissue-adapted CD4^+^ T-cells.

Finally, to determine whether CD49a expression marks functionally distinct CD4^+^ and CD8^+^ T_RM_ populations, cytokine responses were examined in CD4^+^CD103^+^ and CD8^+^CD103^+^ T-cells after anti-CD3/CD28 stimulation. CD4^+^CD103^+^ T-cells showed limited differences in polyfunctional cytokine profiles between CD49a^+^ and CD49a^−^ subsets, except for IL17a, which was slightly higher in CD49a^−^ cells (Supplemental Figure 4A). In contrast, CD8^+^CD103^+^ T-cells showed a significantly increased IFNγ-TNF polyfunctional response (p<0.0001) in CD49a^+^ compared with CD49a^−^ (Supplemental Figure 4B). These findings align with similar studies of skin T_RM_ cells (33) and identify CD49a expression as a marker for functionally distinct CD8^+^CD103^+^ T cells enriched for type 1 inflammatory cytokine production.

### Circulating CD4^+^ T-cells undergo divergent tissue-adaptive programs upon entry into the inflamed vitreous

Having established the expression of canonical T_RM_ markers in vitreous-infiltrating CD4^+^ and CD8^+^ T-cells, we next sought to define the transcriptional programs underlying local tissue adaptation. This question is particularly relevant in PSU, given that the normal vitreous is largely devoid of immune cells, implying that all infiltrating populations arise from the circulation and undergo local tissue adaptation. Also, the current definition of tissue residency has expanded beyond surface marker expression to include distinct transcriptional and epigenetic programs that define T_RM_ identity and function (9,12). To resolve this, we performed scRNA-Seq and TCR-Seq on paired blood and vitreous samples from patients with chronic uveitis. Four patients with chronic PSU (all NIU patients) and severe vitreous haze, who required therapeutic vitrectomy for non-resolving inflammation, were included in the study. None of these patients had any known systemic inflammatory disease. Paired peripheral blood and vitreous samples were collected, and total immune cells were processed for 5’-scRNA and TCR-Seq on the 10x Genomics platform. Following standard quality control, normalization, and integration of the paired blood–vitreous datasets, CD4^+^ and CD8^+^ T-cells were independently clustered using an unsupervised approach. Cluster-wise differential expression and clonotype analyses were then performed to identify the tissue-associated transcriptional states. Reference-based annotation using publicly available atlases (for example, Azimuth) (36) did not adequately capture vitreous T-cell clusters as current reference datasets lack ocular tissue–specific inflammation-adapted T-cell states. Hence, cluster identities were assigned through an integrated assessment of transcriptional profiles (Figure 4, A-C, Supplementary Table 1), clonal expansion (Figure 4D-E), and differential gene expression (Figure 4F, Supplementary Figure 5A-G).

**FIGURE 4:**
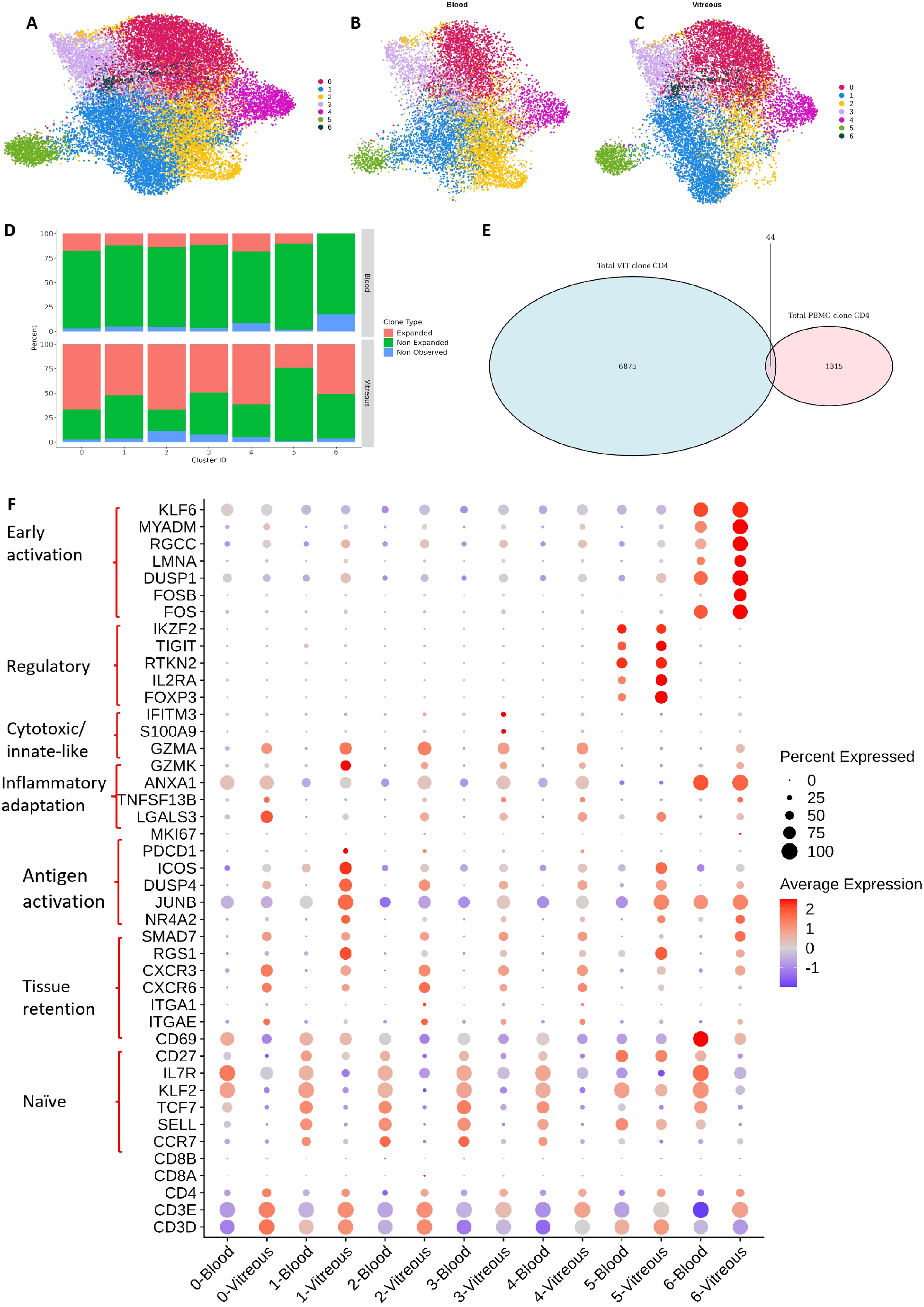
Circulating CD4^+^ T-cells undergo divergent tissue-adaptive programs upon entry into the inflamed vitreous. **A**. Combined UMAP visualization of unsupervised clustering of pooled blood and vitreous samples revealed 7 distinct CD4^+^ T cell populations from four patients with chronic posterior segment uveitis (n = 21,746). **B-C**. UMAP visualization of unsupervised clustering revealed 7 distinct CD4^+^ T cell populations in blood (n = 9589) and vitreous (n = 12,157) samples, respectively. **D**. Distribution of clonal expansion across CD4^+^ T cell clusters in blood (top) and vitreous (bottom), showing a higher proportion of expanded clones within specific vitreous clusters compared to blood. Clone types are categorized as expanded, non-expanded, or not observed. **E**. Overlap of TCR clonotypes between blood and vitreous CD4^+^ T-cells demonstrating minimal sharing, with the majority of vitreous clonotypes being compartment-specific, indicating local clonal expansion and/or selective retention. F. Dot-plot showing expression of key gene signatures across clusters in blood and vitreous, grouped by different functional categories. Dot size represents the percentage of cells expressing the gene, and color intensity reflects the average expression level. Scaled data (Z-score transformation), which centers each gene’s expression around a mean of 0, were used.

To define the full spectrum of CD4^+^ T-cell states in circulation and tissue, we first analyzed transcriptional profiles of clonotypes unique to the blood and vitreous compartments, respectively. This approach was adopted as most clones were unique to the respective compartments. It captures the maximal phenotypic diversity within each compartment, even though with a bias toward terminal or compartment-enriched states. Differential gene expression from the scRNA-Seq data resolved the CD4^+^ cells into seven different subsets (Supplemental Figure 6 A). To interpret these clusters biologically, we adopted a focused panel of markers representing key functional programs, from cluster-specific differentially expressed genes, and established T-cell literature. Circulating CD4^+^ T-cells expressed naïve/central memory markers (*CCR7, TCF7, SELL, KLF2, CD27*, and *IL7R*) across nearly all clusters. No major differences were noted between the clusters except for the expression of regulatory markers (*FOXP3, IL2RA, RTKN2, TIGIT, CTLA4*, and *IKZF2*) in cluster 5, and immediate-early genes (*FOS, DUSP1, LMNA, RGCC, MYADM*, and *KLF6*) in cluster 6. In contrast, vitreous-infiltrating CD4^+^ T cells displayed multiple tissue-adapted phenotypes. Tissue-retention markers were expressed in all clusters, though the patterns differed between clusters. *CXCR3* appeared to be the most ubiquitous, being expressed in all vitreous clusters. *CXCR6* and *ITGAE* (CD103) were expressed primarily in clusters 0 and 2, though *CXCR6* was also expressed in clusters 1, 3, and 4. *CD69* expression was paradoxically low across all vitreous clusters but high in one blood cluster (cluster 6). This could be attributed to the short half-life (<60 minutes) of *CD69* mRNA in activated T-cells (37). To validate this hypothesis, we compared the median fluorescence intensity (MFI) of CD69 in blood and vitreous CD4^+^ cells from our earlier experiments, which revealed higher MFI in the vitreous across all CD69 compartments (Supplemental Figure 7 A-B). The naïve T cell markers (*CCR7, TCF7, SELL, KLF2, CD27*, and *IL7R*) were uniformly downregulated in all clusters except the regulatory cluster 5, where *SELL, KLF2*, and *CD27* were upregulated. The downregulation of *KLF2* is particularly remarkable, given that it promotes the expression of several key trafficking receptors, notably *S1PR1* (38).

Among the individual clusters, cluster 1, expressing *NR4A2, JUNB, DUSP4, ICOS, PDCD1*, and *GZMK*, suggested chronic antigenic stimulation with adaptive restraint (39). While *GZMK* is more commonly associated with CD8^+^ T-cells, its expression in CD4^+^ cells marks a distinct pro-inflammatory lineage (THK) distinct from classical Th1, Th2, or Th17 cells (40). Clusters 3 and 4 also expressed *CXCR3* and *CXCR6*, consistent with their T_RM_ identity, with cluster 3 additionally expressing the cytotoxicity marker *GZMA* (Supplementary Figure 5). Cluster 5 was a *FOXP3*^+^ regulatory T-cell (Treg) population also expressing *IL2RA, RTKN2, TIGIT*, and *IKZF2*. The coexpression of *IKZF2* (Helios), *TIGIT*, and *IL2RA* (CD25) in Tregs identifies a highly suppressive, stable, and “activated” subset in the context of a tumor microenvironment (41). The regulatory marker, *RGS1*, was upregulated in vitreous clusters 1 and 5. While extensively studied for CD8^+^ cells, *RGS1* has also been reported in CD4^+^ T_RM_ and Treg cells in barrier tissues (42). Most clusters were enriched for expanded clonotypes in the vitreous, particularly Clusters 0 and 1, supporting antigen-driven proliferation and local persistence (Figure 4D). This was further reinforced by the consistent expression of *HLA-DR* across clusters 0 to 4 (Supplementary Figure 5). However, the *FOXP3*^+^ regulatory cluster (Cluster 5) showed limited clonal expansion, suggesting that these cells are likely polyclonal rather than antigen-specific. Together with the upregulation of *SELL, KLF2*, and *CD27*, these findings suggest that the vitreous Tregs assume migratory or less differentiated states, being recruited directly from the circulation and/or through local induction within the vitreous microenvironment. These also appear consistent with experimental observations of Foxp3^+^ functionally competent Tregs derived from the circulation that were noted in the vitreous following adaptive transfer of IRBP-specific T cells in mice (43,44). Finally, cluster 6 was characterized by high expression of immediate-early genes (*FOS, FOSB, DUSP1, LMNA, RGCC, MYADM*, and *KLF6*), consistent with a transient activation or a stress-response state rather than a stable lineage (45). Collectively, these findings indicate that vitreous CD4^+^ T-cells comprise a heterogeneous mixture of differentiated effector, regulatory, and tissue-resident populations that are largely distinct from their circulating counterparts.

To directly assess tissue adaptation, we next examined clonotypes shared between blood and vitreous, enabling within-clone comparisons across compartments (Figure 4E, Supplementary Table 1). Despite limited overlap in CD4^+^ clonotypes, shared clones provided critical mechanistic insight into tissue-induced transcriptional changes. Across multiple clusters, shared clonotypes consistently downregulated circulation-associated genes (*KLF2, SELL, TCF7*). Clusters 2 and 3 were notable for high expression of cytolytic genes (*GNLY, FGFBP2, NKG7*) in the blood, suggesting possible egress from the vitreous into the peripheral blood. This was further supported by upregulation of *S1PR5* and *CX3CR1* in cluster 3 blood cells. The corresponding vitreous cells (cluster 2) were marked by hyper-transcription of specific TCR chains (*TRBV2, TRAV23DV6*), suggesting intense local antigen stimulation and massive clonal expansion. Cluster 0 in blood showed a Th-17 bias (*CCR6, TAGAP*), while in the vitreous, it showed localized clonal expansion (*TRBV4-2*), and expression of drivers of B-cell survival (*TNFSF13B*) (46). The vitreous cluster 1 shared clones also had supporting markers for a B-cell niche (*GPR183*), though most markers suggested chronic antigenic stimulation (*PDCD1, CTLA4, TOX, ICOS, RGS1*). Thus, the shared clonotype analysis reveals that when circulating CD4^+^ T-cells enter the vitreous, they adopt varied transcriptional programs, ranging from effector to cytotoxic and tissue-resident states, further reinforcing the functional plasticity of CD4^+^ T-cells within the ocular microenvironment.

### CD8^+^ T-cells undergo selective tissue adaptation within the vitreous microenvironment

In contrast to the CD4^+^ population, analysis of unique clonotypes in the blood and vitreous revealed that the CD8^+^ cells formed more distinct transcriptional clusters in the blood than in the vitreous (Supplemental Figure 8A). As for CD4^+^ cells, we combined transcriptional profiles (Figure 5, A-C, Supplementary Table 1), clonal expansion (Figure 5D-E), and differential gene expression (Figure 5F, Supplementary Figure 9 A-G) for the cluster annotation. Overall, the CD8^+^ T-cells resolved into eight distinct clusters lineage. In the blood, the CD8^+^ cells segregated into multiple cytotoxic effector populations (clusters 3, 5, and 6) expressing *GNLY, GZMB, PRF1*, and *NKG7*, and a naïve-like population (cluster 7) characterized by *CCR7, SELL, TCF7, LEF1*, and *IL7R*. Cytotoxic markers were also expressed in the remaining clusters, except in clusters 2 and 7. Additional markers, such as *KLF2* and *IL7R*, were broadly expressed, consistent with circulating T-cell phenotypes. A distinct HLA class II–expressing population (cluster 5), marked by *HLA-DRA, HLA-DRB1*, and *CD74*, was also present in the circulation, suggesting that this phenotype is not restricted to the tissue environment In contrast to the tissue diversification in the CD4+ compartment, the vitreous CD8^+^ T cells had broadly similar transcriptional profiles to those in blood. Nearly all clusters expressed markers of chronic antigen exposure, including *DUSP4, JUNB, NR4A2*, and *ICOS. GZMK*, a marker of cytokine-driven activation (47), was also broadly expressed, except in clusters 3 and 7. Conversely, expression of cytotoxicity markers was attenuated in all clusters, except cluster 3, which was enriched in *GNLY*. As expected, the tissue retention markers, primarily *CXCR3* and *RGS1*, were expressed in all clusters except cluster 7. *ZNF683* (Hobit) was enriched only in clusters 3 and 7, while the remaining markers – *CXCR6, ITGAE*, and *ITGA1*-were modestly expressed across multiple clusters. As with the CD4^+^ cells, CD69 expression was downregulated in most vitreous clusters, except 0 and 7 (32). Cluster 5 was defined by robust expression of antigen presentation machinery, including *HLA-DRA, HLA-DRB1*, and *CD74*, indicating the presence of an unconventional HLA class II–expressing CD8^+^ population that may contribute to local immune amplification. These cells have been identified in specific clinical contexts, such as chronic viral infections and drug hypersensitivities (48,49). Cluster 6 was enriched for immediate-early and stress-response genes (*FOS, FOSB, DUSP1, LMNA, MYADM*, and KLF6) (45), as well as the cell-cycle regulator *RGCC* (50). Notably, cluster 7 was enriched for *TRDV1-* and *TRDC*-expressing CD8^+^ cells. Such NK-like CD8^+^ cells, described in TB and chronic inflammatory conditions, were found to be hyporesponsive to TCR-mediated signaling yet capable of producing CD16-mediated cytolytic responses (51). This cluster also expressed *JUNB* and *ZNF683*, highlighting its robust tissue adaptation.

**FIGURE 5:**
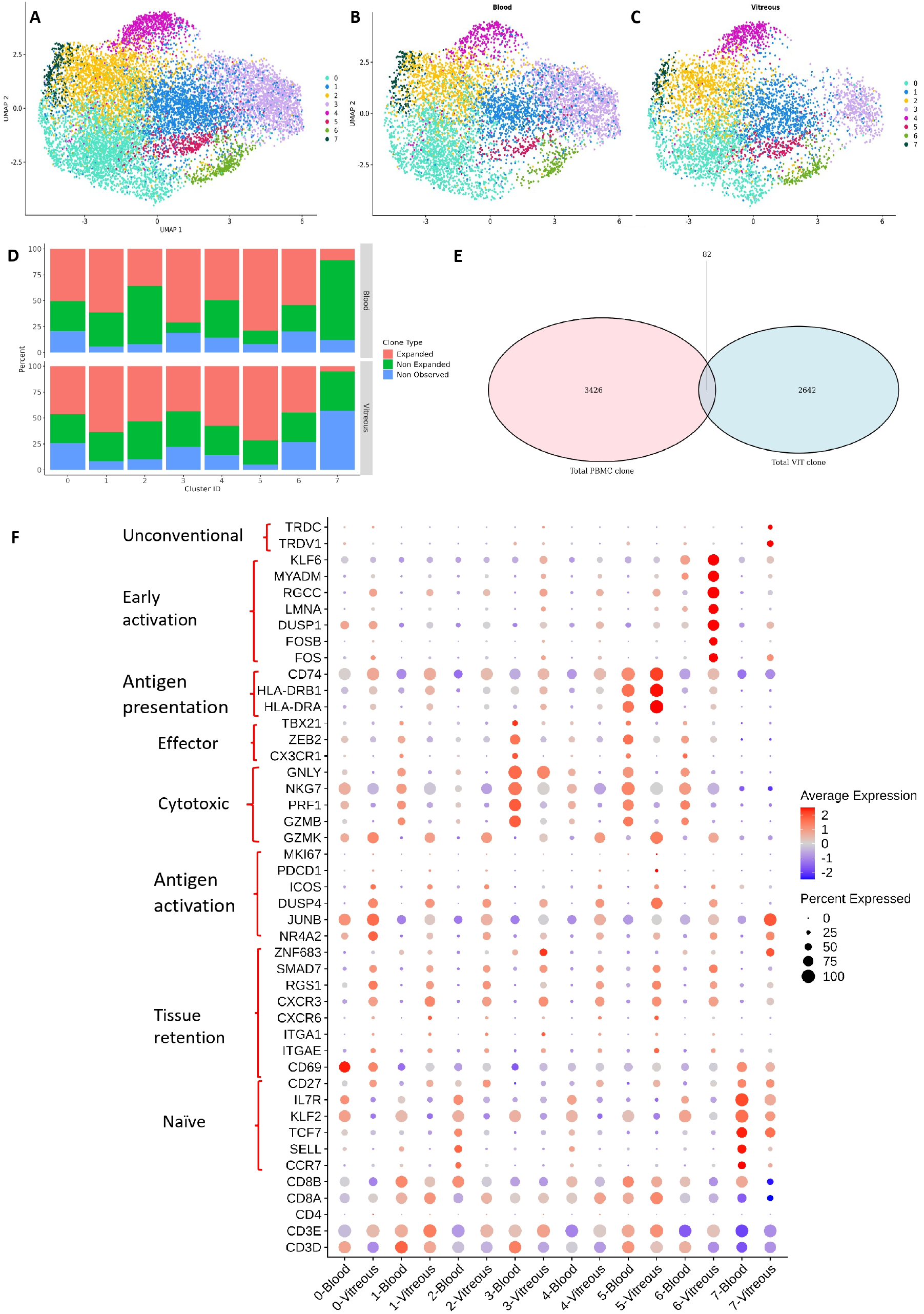
CD8^+^ T-cells undergo selective tissue adaptation within the vitreous microenvironment. **A**. Combined UMAP visualization of unsupervised clustering of pooled blood and vitreous samples revealed 8 distinct CD8^+^ T cell populations from four patients with chronic posterior segment uveitis (n = 11,642). **B-C**. UMAP visualization of unsupervised clustering revealed 8 distinct CD8^+^ T cell populations in blood (n = 6448) and vitreous (n = 5194) samples, respectively. **D**. Distribution of clonal expansion across CD4^+^ T cell clusters in blood (top) and vitreous (bottom), showing a higher proportion of expanded clones within specific vitreous clusters compared to blood. Clone types are categorized as expanded, non-expanded, or not observed. **E**. Overlap of TCR clonotypes between blood and vitreous CD8^+^ T-cells demonstrating minimal sharing, with the majority of vitreous clonotypes being compartment-specific, indicating local clonal expansion and/or selective retention. F. Dot-plot showing expression of key gene signatures across clusters in blood and vitreous, grouped by different functional categories. Dot size represents the percentage of cells expressing the gene, and color intensity reflects the average expression level. Scaled data (Z-score transformation), which centers each gene’s expression around a mean of 0, were used.

Clonal analysis further supported a model of systemic priming followed by tissue recruitment of CD8^+^ T cells (Figure 5E, Supplementary Table 1). Clonal expansion was consistently greater in the blood across clusters, with expanded clones comprising a larger fraction of circulating CD8^+^ T cells, whereas vitreous populations (except cluster 2) either mirrored or showed less clonal expansion than their corresponding peripheral blood clusters. There was greater clonal sharing between blood and vitreous compartments than for CD4^+^ cells, indicating that many tissue-infiltrating CD8^+^ T cells are derived from circulating clones. Consistent with this, differential expression analysis of the shared clones demonstrated a conserved program of tissue adaptation, with vitreous CD8 T-cells upregulating regulatory stress-response genes (*FOS, JUN, DUSP1/2/4, RGCC*) in multiple clusters, with limited expression of tissue residency markers *RGS1* and *ICOS* (cluster 0). *PMEPA1*, a negative regulator of TGF-ß signaling, was enriched in clusters 0, 1, 2, and 5, showing that these cells were modulating their sensitivity to the TGF-ß-rich ocular microenvironment. Collectively, these data support a model in which CD8^+^ T-cells in uveitis are systemically expanded and dynamically recruited into the vitreous, where they undergo regulatory reprogramming without establishing a uniform tissue-resident phenotype. In the blood compartment, *CX3CR1* upregulation was observed in clusters 1 and 2, indicating their egress from the vitreous. As with the CD4^+^ cells, high expression of cytolytic genes (*GNLY, GZMB*) was noted in clusters 2 and 3. However, in the total CD8^+^ population (not shared clones alone), only cluster 3 was highly expanded, and upregulated all cytolytic markers in the blood (Figure 5D, F). Taken together, the shared clones of cluster 2 could potentially represent recirculating CD8^+^ cells from the vitreous that had acquired a cytolytic phenotype on entering the vitreous.

### Independent vitreous cohort confirms heterogeneous transcriptional programs in CD4^+^ and CD8^+^ T-cells

To assess the reproducibility of T-cell states identified in the exploratory cohort, we analyzed vitreous samples from an independent validation cohort (chronic PSU) and performed parallel clustering of CD4^+^ and CD8^+^ T cells. Clusters were identified with canonical marker genes and then compared across cohorts using both gene-level overlap (Jaccard similarity index) (Figure 6A-B) and visualization of marker expression patterns by the respective dot plots (Figure 6C-D).

**FIGURE 6.**
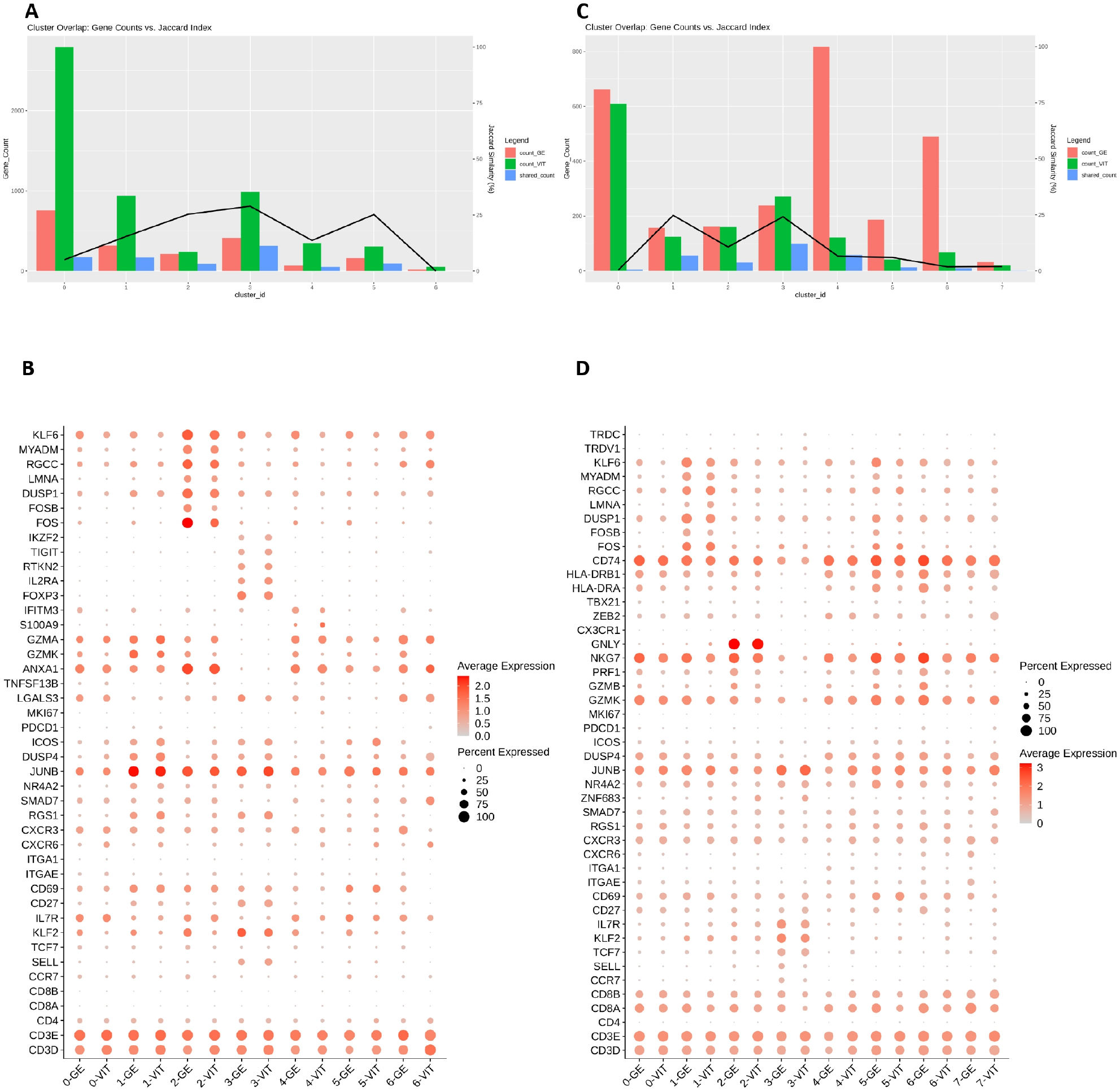
Independent vitreous cohort confirms heterogeneous transcriptional programs in CD4^+^and CD8^+^ T-cells. (A) Gene-level overlap between exploratory (VIT) and validation (GE) cohorts for CD4^+^ T cell clusters. Bar plots represent the numbers of differentially expressed genes identified in each cohort and the number of shared genes per cluster, while the overlaid line indicates Jaccard similarity (%). (B) Gene-level overlap among CD8^+^ T cell clusters across cohorts, as shown in (A). (C) Dot plot showing expression of selected canonical marker genes across CD4^+^ T cell clusters in exploratory (VIT) and validation (GE) cohorts. (D) Dot plot of marker gene expression across CD8^+^ T cell clusters in both cohorts. Dot size represents the percentage of cells expressing each gene, and color intensity indicates average expression.

Among CD4^+^ T-cells, several clusters (2, 3, and 5) demonstrated strong cross-cohort concordance. These clusters showed relatively higher (≥25%) Jaccard similarity (Figure 6A) and highly consistent marker expression patterns (Figure 6B) across cohorts. Chronic antigenic stimulation, as defined by *NR4A2, DUSP4, JUNB, RGS1, ICOS*, and *GZMK*, was consistently observed in multiple clusters, but most prominently in cluster 1. A distinct population characterized by high expression of immediate-early genes (*FOS, FOSB, DUSP1, LMNA, RGCC, MYADM*, and *KLF6*), was noted in cluster 2. A canonical *FOXP3*^+^*IL2RA*^+^*IKZF2*^+^i regulatory T cell compartment was robustly reproduced in cluster 3. The remaining clusters demonstrated lower gene-level overlap, likely reflecting their dynamic transcriptional nature. Nevertheless, core lineage-defining markers remained conserved, supporting the preservation of underlying biological states despite variability in individual gene lists.

CD8^+^ T-cells exhibited a different pattern, with generally lower Jaccard similarity (Figure 7C), across clusters despite clear conservation of key transcriptional programs (Figure 7D). Activated states characterized by immediate-early genes (FOS, FOSB JUN, DUSP family) were found in cluster 1, and naïve/central memory markers (*CCR7, SELL, TCF7, KLF2* and *IL7R*) in cluster 3 of both cohorts, although with variable gene composition. Cytotoxic populations expressing *GZMB, PRF1, GNLY*, and *NKG7* were more prominent in the validation cohort (clusters 2, 5, and 6). Hybrid populations expressing HLA class II molecules were found in both clusters 5 and 6, again more prominently in the validation cohort. These findings indicate that CD8^+^ T-cells in the inflamed vitreous exist along a continuum of overlapping functional programs, leading to reduced gene-level concordance compared with CD4^+^ T-cells.

Together, these analyses demonstrate that, despite the expected variability in individual differentially expressed genes, the major CD4^+^ and CD8^+^ T-cell states identified in the exploratory cohort are reproducibly observed in an independent validation cohort. The consistent preservation of key transcriptional programs across datasets supports the robustness of our clustering and annotation strategy and validates the core immune architecture in chronic PSU.

## Discussion

In this study, we provide a comprehensive multimodal characterization of tissue-adapted T-cell responses in the vitreous of patients with chronic PSU who underwent vitrectomy (vitreous biopsy) for diagnostic and/or therapeutic indications. We integrated high-dimensional phenotyping and antigen-specific functional assays with paired single-cell transcriptomic and TCR repertoire analyses. By directly sampling the vitreous, a compartment that faithfully reflects the ocular inflammatory microenvironment, we examined how circulating T cells are transcriptionally reprogrammed within the human eye. The transcriptional programs identified in both CD4^+^ and CD8^+^ T-cells were also independently validated in a separate cohort of PSU patients. Our findings establish that chronic uveitis is sustained by heterogeneous, tissue-adapted T-cell populations that extend beyond canonical T_RM_ marker expression, with CD4^+^ and CD8^+^ T-cells exhibiting distinct patterns of tissue adaptation within the ocular microenvironment.

Our understanding of intraocular immune responses in chronic uveitis has been constrained by the generally self-limited nature of experimental autoimmune uveitis (EAU). Inflammation in conventional EAU has been shown to peak within 2–3 weeks after immunization with retinal autoantigens and then resolve spontaneously (52,53). However, chronic persistent inflammation associated with angiogenesis, has been noted up to 120 days following immunization of C57BL6 mice with IRBP (peptide 1-20) (54). In this model, CD8^+^CD103^+^ cells were found in the retina during the persistent phase (days 36-43) of inflammation (55). Recent modifications to the immunization protocol have prolonged the duration of inflammation (56), and identified long-lived CD44^hi^IL-7R^+^IL-15R^+^CD4^+^ memory T-cells in the eye up to 3 months post-immunization (57). Other hallmarks of tissue adaptation were not reported in these studies. More recently, CD8^+^CD103^+^ cells were identified in both non-inflamed and inflamed uveal tissue from human donor eyes, and in the mouse anterior uvea after the resolution of EAU (58). In the same study, paired scRNA and TCR-Seq of aqueous samples from anterior uveitis found *ITGAE, ITGA1*, and *CXCR6* expression in the expanded T-cell population. Another study of the healthy mouse anterior uvea revealed CD3^+^CD4^−^CD8^−^CD69^+^γδTCR^+^IL-23R^+^ tissue-resident cells capable of driving uveitis-like inflammation upon local IL-23 activation (59). Other single-cell studies of human aqueous samples (60-62), or EAU (63), have not reported clear T_RM_-associated programs. Together, these observations suggest that tissue adaptation occurs within the eye but remains incompletely defined. Our study extends these findings by identifying the vitreous body as a site of sustained immune adaptation, where long-lived T-cells acquire tissue-resident features that distinguish them from their circulating counterparts.

Unlike circulating fluids such as aqueous humor or CSF, the vitreous possesses a defined three-dimensional architecture stabilized by an organized collagen-hyaluronan matrix (18). This matrix is actively maintained by hyalocytes and creates a tissue microenvironment capable of shaping local immune responses. However, the pattern of tissue adaptation in the vitreous differs from that in classical barrier tissues such as the skin, gut, and lung. In these tissues, T_RM_ maintenance depends heavily on epithelial anchoring through CD103- and CD49a-mediated interactions (7-13). In the vitreous too, CD8^+^ T-cells preferentially expressed these retention-associated markers, but they remained clonally and phenotypically closer to blood CD8^+^ T cells in gene expression. In contrast, CD4^+^ T-cells showed greater enrichment of the homing receptor CXCR6, stronger antigen-specific responses, localized clonal expansion, and compartment-specific transcriptional reprogramming. Vitreous T-cells also upregulated *NR4A2, DUSP4, JUNB, ICOS, PDCD1*, and *GZMK*. These molecules may reflect adaptive restraint programs induced by chronic antigen exposure and local immunomodulatory factors (64). These may include cellular elements (retinal Müller glia, and retinal pigment epithelium) (65, 66), and soluble factors (TGF-β, retinoic acid, and others) (64, 67), although these were not directly interrogated in this study. Together, these observations suggest that tissue adaptation within the vitreous is not a single resident-memory state but a collection of adaptation programs, including structural retention and engagement with the inflammatory niche, that are differentially utilized by CD8^+^ and CD4^+^ T-cells.

A striking feature of the vitreous CD4^+^ compartment was the coexistence of marked local adaptation and evidence of ongoing cellular plasticity. In barrier tissues, CD4^+^ T_RM_ cells are generally considered less terminally differentiated and more prone to recirculation than their CD8^+^ counterparts (68). Likewise, the vitreous CD4^+^ T-cells did not converge into a single dominant resident phenotype but instead segregated into multiple clonally expanded tissue-adaptive states with distinct functional and transcriptional programs. Although antigen-specific and polyfunctional responses were enriched within CD69^+^CD103^+^CD4^+^ cells, only a subset of these cells exhibited robust effector function. Together with the expression of *NR4A2, DUSP4, JUNB, ICOS, PDCD1*, and *GZMK*, this observation suggests that chronic antigen exposure in the immune-privileged vitreous induces an adaptive restraint on effector functions (69,70). Such programs may limit excessive tissue damage while preserving local immune persistence. Additional evidence for selective pressures within the vitreous came from the Treg compartment, which remained largely polyclonal and distinct from the expanded effector populations (71). Collectively, our data suggest that tissue-adapted CD4^+^ T cells in chronic uveitis exist along a spectrum of states whose inflammatory potential is continuously shaped by the local immune environment.

In contrast to the diversity of the CD4^+^ compartment, vitreous CD8^+^ T cells exhibited a more uniform pattern of tissue adaptation. They showed substantial clonal overlap with the circulation, with relatively limited compartment-specific reprogramming, despite expressing higher levels of the retention-associated markers CD103 and CD49a. This apparent paradox may reflect a fundamental difference in how the two lineages adapt to tissue environments. Whereas CD4^+^ cells occupied multiple tissue-adaptive states, CD8^+^ cells appeared to converge on a more stable phenotype characterized by reduced expression of cytotoxic effector molecules and increased expression of regulatory genes, including *NR4A2, DUSP4, JUNB*, and *RGS1*. Similar restrained CD8^+^ states have been described in chronically inflamed tissues, where persistent antigen exposure and local regulatory signals limit tissue injury without eliminating immune function (72,73). The accumulation of CD8^+^ cells during the chronic phases of experimental and human uveitis (54-58), may therefore reflect prolonged tissue persistence rather than a functional diversification. Collectively, these findings suggest that CD8^+^ T-cells are defined more by long-term tissue persistence than by dynamic adaptation within the vitreous.

Our findings suggest that chronic uveitis may be sustained by tissue-adapted immune states generated by the vitreous microenvironment and are therefore incompletely targeted by therapies directed primarily at circulating immunity. Similar mechanisms have been implicated in chronic inflammatory diseases of the skin, gut, kidney, and central nervous system, where T_RM_ cells contribute to persistent inflammation despite systemic immune control (74-77). The distinct patterns of adaptation observed in vitreous CD4^+^ and CD8^+^ T-cells further suggest that pathogenic tissue immune states extend beyond canonical T_RM_ marker expression. These observations highlight the importance of defining tissue-adapted immune states within target organs and support the development of therapeutic approaches aimed at disrupting local retention and other mechanisms that sustain chronic intraocular inflammation (78-79).

Several limitations should be considered while interpreting this study. As in any human study, the clinical cohort was heterogeneous with differences in disease etiology, duration, and treatment status. While it could influence the T-cell composition and functional state, it also reflects the real-world complexity of chronic posterior segment uveitis. Secondly, we found partial discordance between flow cytometry-based phenotyping and scRNA-Seq-based transcript detection. This is also a biological reality, as demonstrated in CITE-Seq studies for select markers such as CD4 (80). Thirdly, although the vitreous body constitutes a tissue-like immune niche, our study could not determine the spatial organization of immune cells within the vitreous matrix or their interactions with adjacent tissue (retina and choroid) compartments. Finally, the cross-sectional design of this study did not allow assessment of the temporal dynamics of the immune response, including any perturbation studies, to formally establish long-term tissue residency in the vitreous. Notwithstanding these limitations, separating classic tissue residency from functional adaptation provides a fresh framework for understanding chronic inflammation, with broad implications for ocular and systemic autoimmune diseases.

## Methods

### Patient selection

The study was approved by the ethics committee of LV Prasad Eye Institute, Hyderabad, India, and followed the tenets of the Declaration of Helsinki. Patients with non-infectious posterior uveitis who received therapeutic or diagnostic pars plana vitrectomy were included in the study following a written, informed consent (Supplementary Table 2). The only exceptions were patients diagnosed with ocular tuberculosis, either before or after inclusion into the study. The majority (85.1%, n=63) of the patients were not on any systemic anti-inflammatory therapy at the time of sampling. Those on systemic anti-inflammatory therapy were included only if they had a worsening of inflammation in the past two weeks. None of the patients included in the paired blood and vitreous sampling were receiving systemic anti-inflammatory therapy or diagnosed with any systemic inflammatory disease.

### Vitreous cell isolation

Vitreous fluid samples were processed as described before (23). Briefly, about 2 cc of undiluted collected in RPMI medium and transported to the laboratory at room temperature. Samples were digested with hyaluronidase for 1 hour at 37°C in a 5% CO_2_incubator to reduce viscosity and facilitate cell recovery. Following digestion, the reaction was quenched with fetal bovine serum (FBS), and samples were diluted with an equal volume of RPMI. Cell suspensions were filtered through a 40 μm cell strainer (Corning) to obtain single-cell suspensions and centrifuged at 600 g for 20 minutes. The supernatant was discarded and cells were washed by centrifugation at 700 g for 7 minutes. The cell pellet was resuspended in complete RPMI medium (RPMI 1640 supplemented with 10% FBS, 100 U/mL penicillin, 100 μg/mL streptomycin, and 2 mM L-glutamine; Gibco/Invitrogen). Cell number and viability were determined using trypan blue exclusion and a hemocytometer. To preserve functional responses, freshly isolated vitreous cells were used for stimulation assays on the same day without cryopreservation.

### Antigen-specific and TCR-mediated stimulation

Vitreous immune cells were evaluated for antigen-specific and TCR-mediated activation. For antigen-specific stimulation, cells (0.75 × 10^5^ cells/mL) were plated in 96-well U-bottom culture plates and stimulated with peptide pools derived from *Mycobacterium tuberculosis* antigens ESAT-6 and CFP-10 (10 μg/mL each; BEI Resources, NR-50711 and NR-50712) or interphotoreceptor retinoid-binding protein (IRBP, peptide 1–20; GenScript), a known uveitogenic antigen. Anti-CD28 antibody (2 μg/mL; Invitrogen) was added to all cultures to provide co-stimulatory signaling. For polyclonal TCR stimulation, cells were stimulated with anti-CD3/CD28–coated beads according to the manufacturer’s instructions. Cells were incubated for 6 hours at 37°C with 5% CO_2_in the presence of Brefeldin A (10 μg/mL) and monensin (2 μM) to enable intracellular cytokine accumulation.

### Flow cytometry

Cells were first stained with fluorochrome-conjugated antibodies against human CD3 (BD Biosciences), CD4 (BD Biosciences), CD8 (BD Biosciences), CD45RA (BD Biosciences), CD62L (BD Biosciences), CD69 (BD Biosciences), CD103 (BD Biosciences), CXCR6 (BD Biosciences), and CD49a (BD Biosciences). Cells were incubated with antibody cocktails for 40 minutes at room temperature in the dark. For intracellular cytokine staining, cells were fixed using Cytofix fixation buffer (BD Biosciences) for 20 minutes at 4°C. Cells were washed with permeabilization buffer and stained intracellularly with antibodies against TNF-α (BD Biosciences), IFN-γ (BD Biosciences), IL-17A (BD Biosciences), and granzyme B (BD Biosciences) for 45 minutes at 4°C in the dark. Cells were washed and resuspended in PBS containing 2% FBS prior to acquisition. Flow cytometry data were acquired on a CytoFLEX S flow cytometer (Beckman Coulter) and analyzed using FlowJo software (v10.10.1).

### Single-cell RNA sequencing and TCR sequencing

Single-cell suspensions were counted and loaded onto the Chromium Controller (10x Genomics) targeting a maximum recovery of 10,000 cells per reaction. Libraries were generated using the Chromium Next GEM Single Cell 5′ V(D)J Reagent Kit (v1.1 chemistry) according to the manufacturer’s instructions. Gene expression and paired TCR libraries were generated using individual Chromium i7 indices. Libraries were sequenced on Illumina platforms to a minimum depth of 20,000 reads per cell for both gene expression and TCR libraries. Raw sequencing reads were processed using Cell Ranger v7.2.0 with the GRCh38 reference genome. Quality control metrics included UMI count per cell, number of detected genes, and mitochondrial gene percentage. Cell filtering was performed using an automated thresholding approach based on boxplot outlier detection. This data-driven approach ensured removal of low-quality cells, empty droplets, and technical artifacts while maximizing retention of biologically relevant cell populations. Doublets were removed using DoubletFinder. Downstream analysis of scRNA-Seq datasets was performed using Seurat v5. Datasets were normalized using SCTransform and highly variable genes were identified using the VST method, selecting the top 3,000 genes for downstream analysis. To remove technical batch effects while preserving biological variation, we integrated multiple samples using Seurat’s canonical correlation analysis (CCA) integration workflow.

CD4^+^ and CD8^+^ T cell populations were initially identified by annotating the single-cell dataset using the Azimuth reference-based annotation tool. Cells predicted to belong to CD4^+^ or CD8^+^ T cell clusters based on Azimuth’s PBMC reference mapping were selected for downstream analysis. To confirm the authenticity of Azimuth-selected cells, marker-based filtering was performed in a hierarchical manner. Cells were first filtered to retain only CD3+ T cells by requiring expression of pan-T cell markers CD3D and CD3E. From the CD3^+^ population, CD4^+^ T cells were selected based on positive expression of the CD4 marker gene. CD8+ T cells were selected based on positive expression of CD8A and CD8B marker genes. Cells co-expressing both CD4 and CD8 markers (CD4^+^CD8^+^ double-positive cells) were filtered out, retaining only CD4^+^CD8^−^ and CD4^−^CD8^+^ populations. This hierarchical marker-based selection strategy allowed us to confidently identify mature CD4^+^ and CD8^+^ T cell populations.

Cluster-specific marker genes were identified using Seurat’s FindAllMarkers() function. Differential expression was assessed using the Wilcoxon rank-sum test comparing each cluster against all other cells. Genes were considered significant cluster markers if they met the following criteria: (1) adjusted p-value < 0.05 using Benjamini-Hochberg correction for multiple testing, (2) positive log_2_(fold-change), indicating enhanced expression in the target cluster relative to other clusters, and (3) minimum percentage of cells expressing the gene > 0.10. Top upregulated markers per cluster were visualized using dot plots to confirm cluster identity and validate biological relevance. To evaluate the reproducibility and translational robustness of the identified T-cell clusters, our discovery dataset was systematically compared against an independent validation cohort using Canonical Correlation Analysis (CCA) in Seurat to mitigate batch effects. Unsupervised clustering was performed on the integrated dataset to define cross-cohort consensus clusters. DEGs were identified for each cluster using the Wilcoxon rank-sum test (p_adj < 0.05). They were segregated into significantly upregulated (positive markers) and downregulated (negative markers) gene sets. The Jaccard Similarity Index was calculated cluster-wise between the two cohorts to formally quantify the transcriptional conservation of these profiles.

### Statistical analysis

Flow cytometry data were analyzed using FlowJo (v10.10.1). Antigen-reactive T-cells were defined as the proportion of CD4^+^ or CD8^+^ T-cells expressing IFN-γ, TNF-α, granzyme B, or IL-17A following stimulation. T cell populations were identified according to the gating strategy shown in Figure X. Statistical analyses were performed using GraphPad Prism (v10.1.0). Data are presented as individual patient values with median and interquartile range. Nonparametric tests were used throughout. Comparisons between multiple unpaired groups were performed using the Kruskal–Walli’s test with Dunn’s post hoc correction. Paired comparisons across multiple subsets were performed using the Friedman test with Dunn’s multiple-comparison correction. P values <0.05 were considered statistically significant.

## Acknowledgments

SB was funded by the DBT Wellcome Trust India Alliance Fellowship in Clinical and Public Health Research (IA/CPHI/18/1/503975) and the Hyderabad Eye Research Foundation.

## Data availability statement

The data that support the findings of this study are available on request from the corresponding author. The data are not publicly available due to privacy or ethical restrictions.

