## Supplemental File for "Distinct Programs of Tissue Adaptation Shape Vitreous CD4^+^ and CD8^+^ T-cell States in Chronic Uveitis"

### Supplementary Figure 1

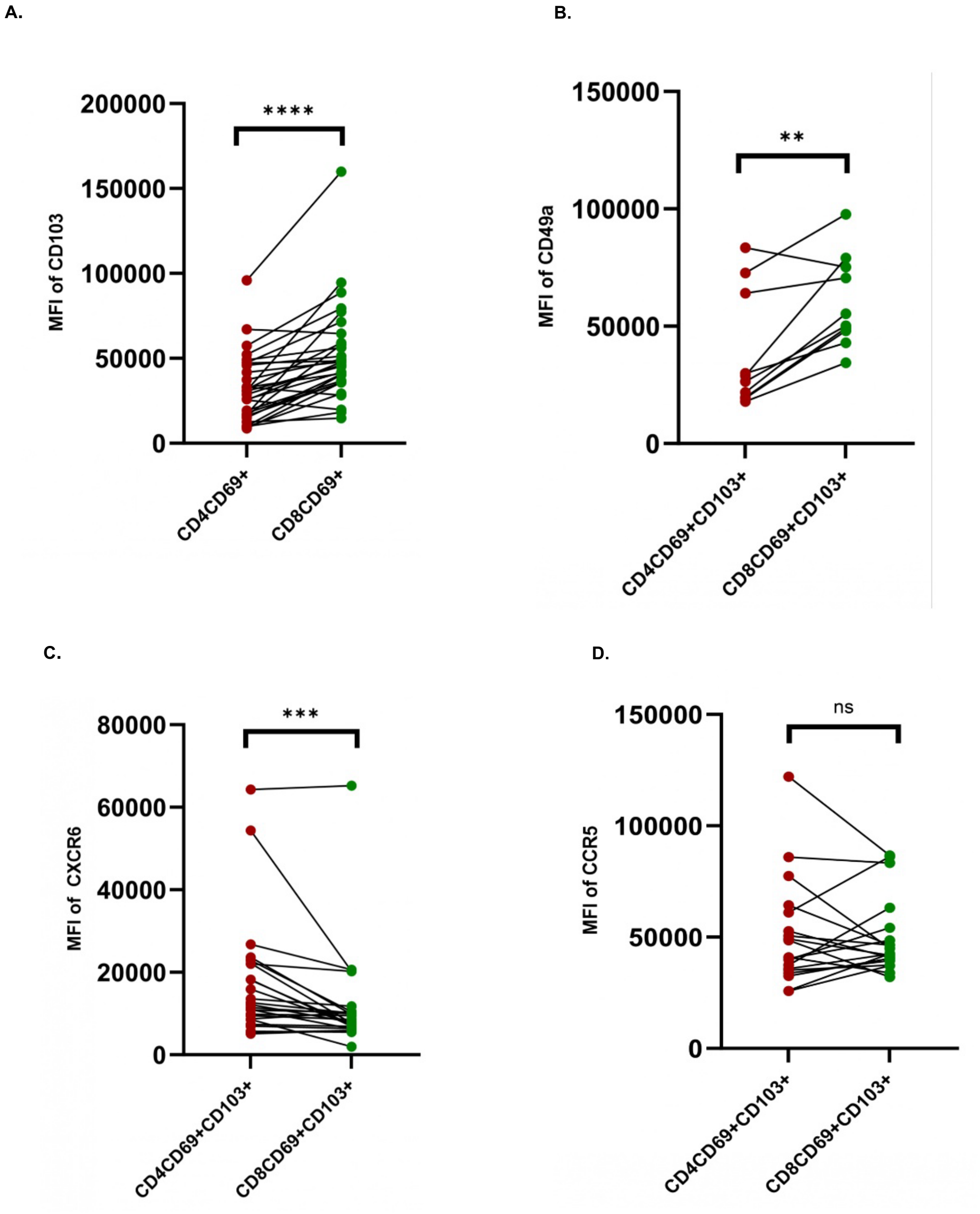


**SUPPLEMENTARY FIGURE 1: A**. Comparison of MFI for CD103 in CD4 and CD8 by paired Wilcoxon sign t test (n=30). **B.** Median fluorescence intensity (MFI) of CD49a expression was assessed in vitreous CD4 and CD8 T cells (n=10). **C.** Median fluorescence intensity (MFI) of CXCR6 (n=25). **D.** Median fluorescence intensity (MFI) of CCR5 (n=19). Statistical significance was determined using a paired Wilcoxon signed-rank test. ***p < 0.001.

Individual *P* values are noted on respective graphs or summarized as: *$P$ < 0.05, **$P$ < 0.01, ***$P$ < 0.001, ****$P$ < 0.0001. In representative histogram images blue peak is represents non stimulated while pink represents the stimulated sample.

### Supplementary Figure 2

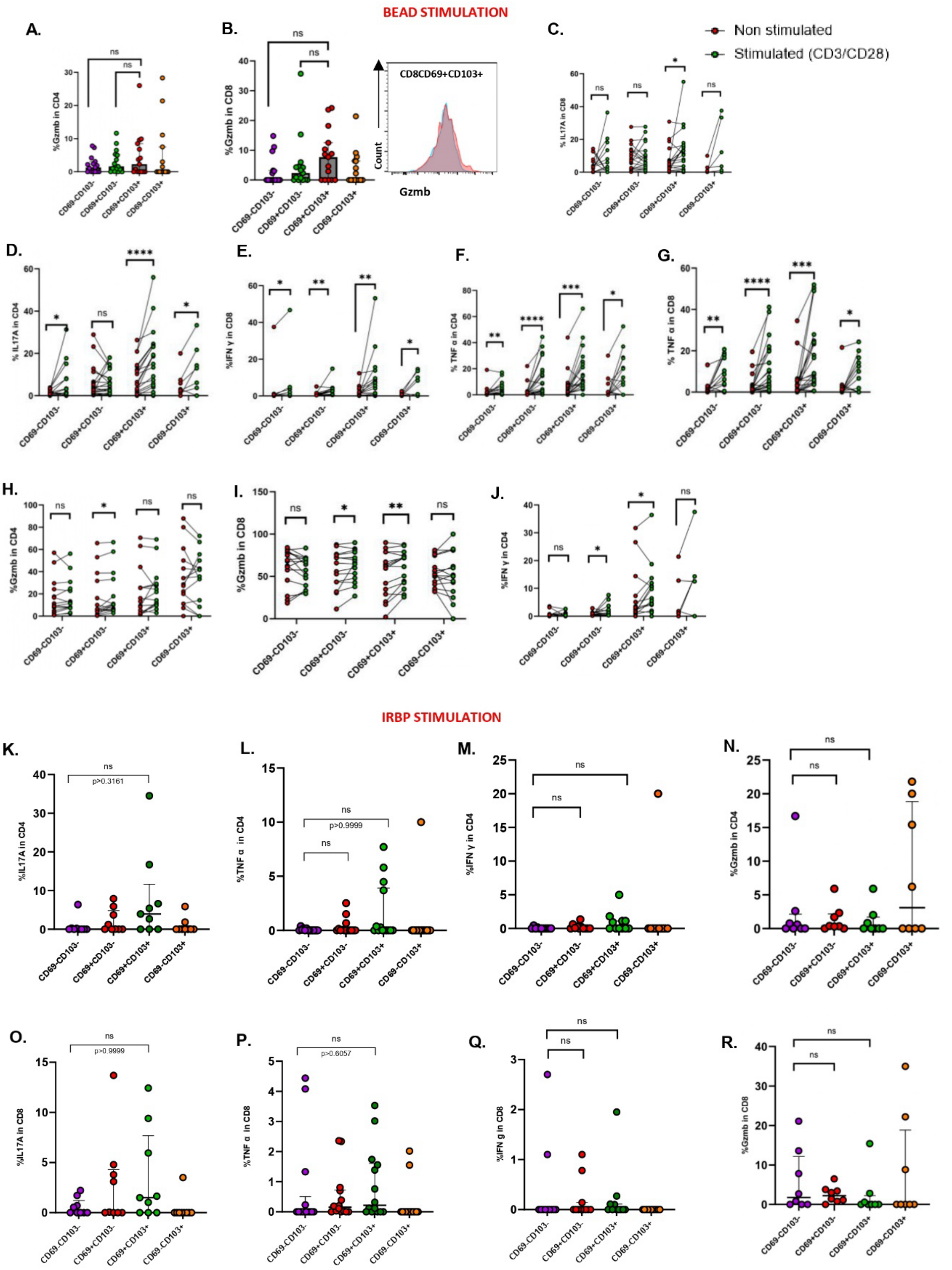


**SUPPLEMENTARY FIGURE 2: A-B.** Monofunctional response of Gzmb in CD4 and CD8 T (n=15) cells upon CD3/CD28 TCR mediated stimulation with representative histogram images. Friedman test for paired data with Dunn's test as post hoc analysis. (**C-J).** Cytokine response of TNFα (n=21), IL-17A (n=21) and IFN-γ (n=15) in CD4 and CD8 T in individual sample upon CD3/CD28 stimulation. Multiple t test (Wilcoxon sign) with Benjamini-Hochberg correction is used for analysis. Each dot represents an individual donor sample, with paired samples connected by lines. **(K-R).** Monofunctional response of TNFα (N=14), IL-17A (n=9), IFN-γ (n=11) and Gzmb (n=12) in CD4 T and CD8 cells upon IRBP stimulation. Friedman test for paired data with Dunn's test as post hoc analysis is used for comparison. Individual *P* values are noted on respective graphs or summarized as: **P* < 0.05, ***P* < 0.01, ****P* < 0.001, *****P* < 0.0001. In representative histogram images blue peak is represents non stimulated while pink represents the stimulated sample.

### Supplementary Figure 3

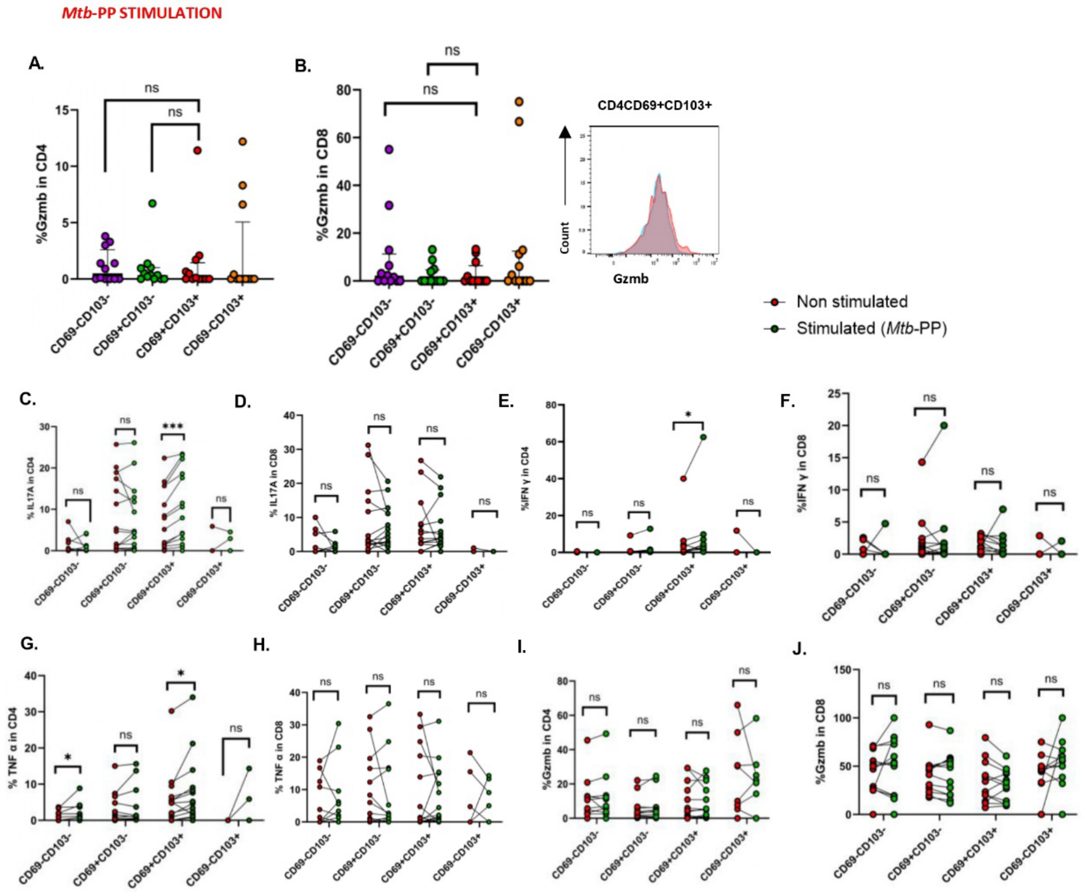


**SUPPLEMENTARY FIGURE 3: A-B.** Monofunctional response of Gzmb in CD4 and CD8 T (n=12) cells upon *Mtb*-PP stimulation with representative histogram images. Friedman test for paired data with Dunn's test as post hoc analysis (**C-J).** Cytokine response of TNFα (n=15), IL-17A (n=15) and IFN-γ (n=11) in CD4 and CD8 T in individual sample upon *Mtb*-PP stimulation. Multiple t test (Wilcoxon sign) with Benjamini-Hochberg correction is used for analysis. Individual *P* values are noted on respective graphs or summarized as: $^*P < 0.05$, $^{**}P < 0.01$, $^{***}P < 0.001$, $^{****}P < 0.0001$. Bars show median ± interquartile ring. In representative histogram images blue peak is represents non stimulated while pink represents the stimulated sample.

### Supplementary Figure 4

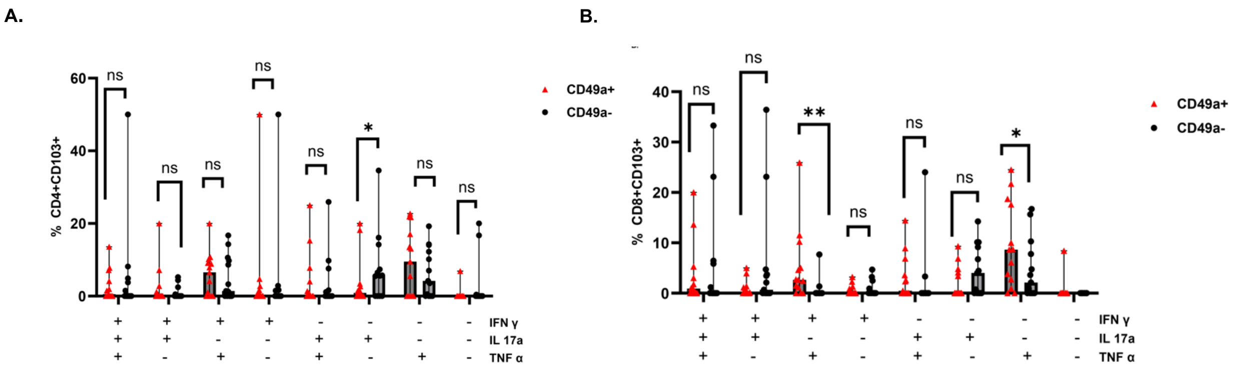


**SUPPLEMENTARY FIGURE 4: CD49a expression modulates the cytokine profile of tissue-residency T cells. A.** Polyfunctional response of TNFα, IL-17A, IFN-γ in CD4CD49a-CD103+ versus CD4CD49a+CD103+ subsets upon CD3/CD28 stimulation (n=13). **B.** Polyfunctional response of TNFα, IL-17A, IFN-γ in CD8CD49a-CD103+ versus CD8CD49a+CD103+ subsets upon CD3/CD28 stimulation (n=13). Multiple t-tests (Wilcoxon signed) with Benjamini-Hochberg correction are done. Individual *P* values are noted on respective graphs or summarized as: $^*P < 0.05$, $^{**}P < 0.01$, $^{***}P < 0.001$, $^{****}P < 0.0001$.

### Supplementary Figure 5

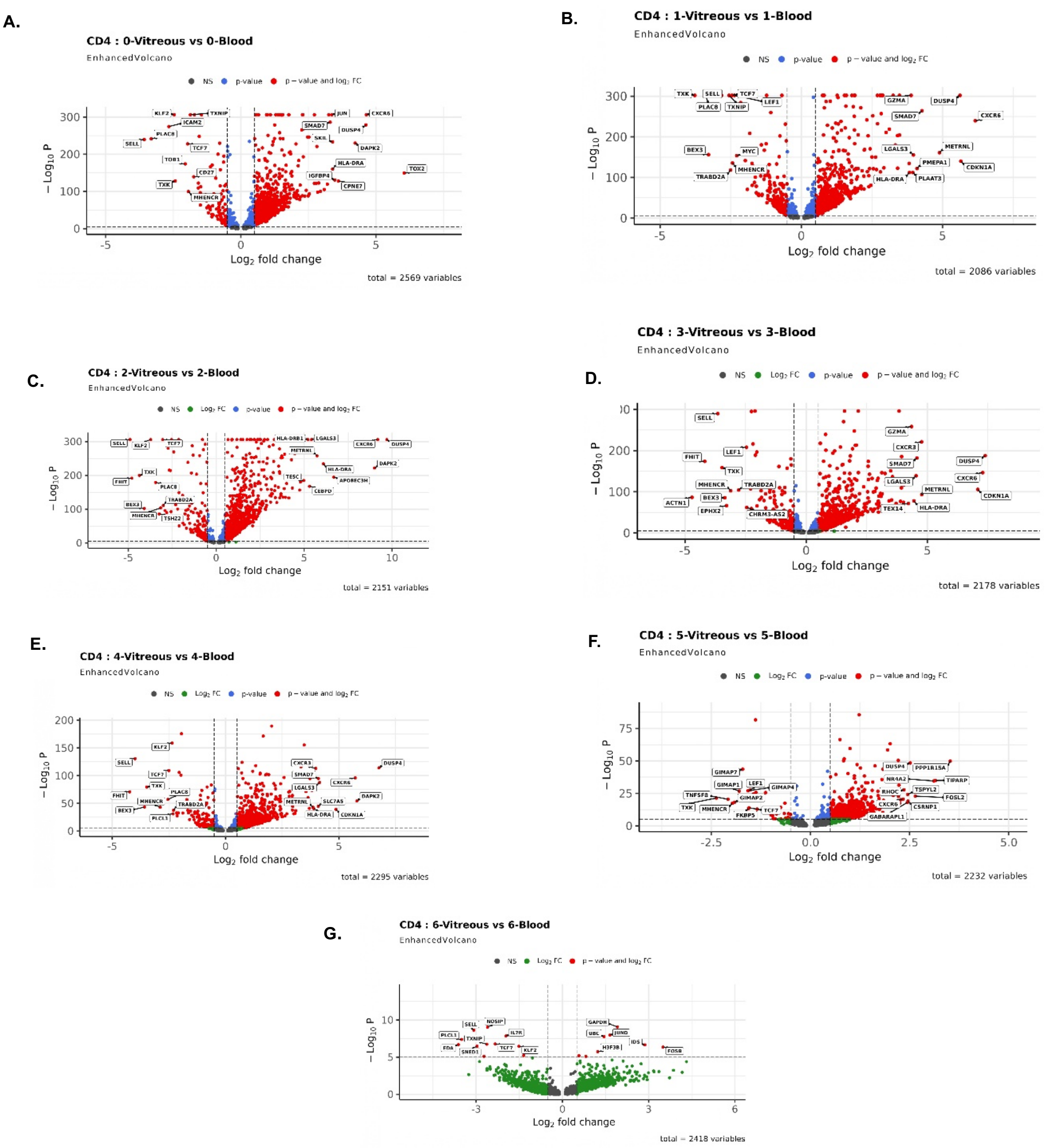


**SUPPLEMENTARY FIGURE 5: A-G** Volcano plots showing differential gene expression between vitreous-derived and peripheral blood CD4$^{+}$ T-cell subsets across multiple clusters. Panels A to G correspond to clusters 0 to 6, respectively. The x-axis represents $\log_2$ fold change (vitreous vs blood), and the y-axis represents $-\log_{10}$(p-value). Each dot represents a gene. Genes not meeting significance thresholds are shown in grey (NS) while genes with significant fold change only are shown in green, genes with significant p-values only are shown in blue, and genes meeting both significance thresholds (adjusted p-value and $\log_2$ fold change) are highlighted in red. Selected differentially expressed genes are annotated. Lower right of each panel indicates total number of genes analysed in each comparison. Dashed vertical and horizontal lines indicate threshold cutoffs for fold change and statistical significance.

### Supplementary Figure 6

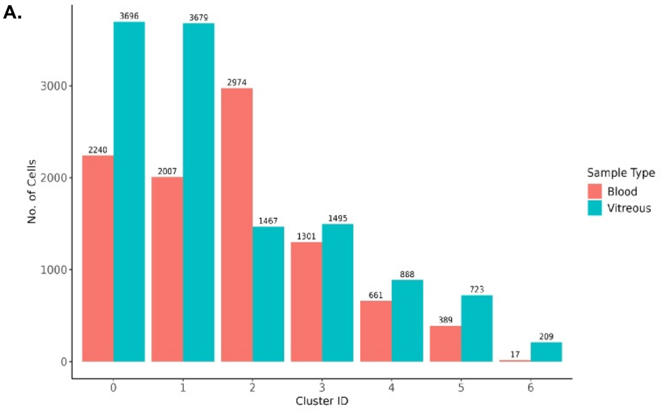


**SUPPLEMENTARY FIGURE 6: A.** Distribution of CD4 T cells across identified clusters in Blood and Vitreous samples. The bar plot shows the number of cells in each cluster (Cluster 0–6) for two sample types: Blood (pink) and Vitreous (blue). Cell counts are indicated above each bar.

### Supplementary Figure 7

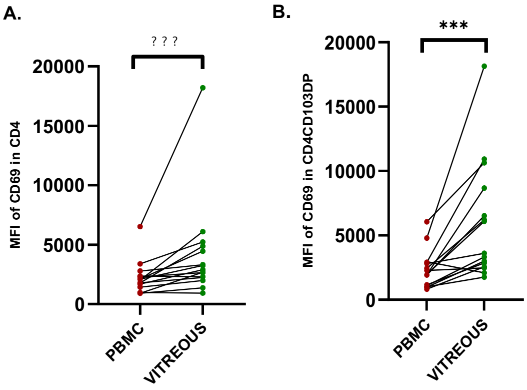


**SUPPLEMENTARY FIGURE 7: A-B.** Median fluorescence intensity (MFI) of CD69 expression was assessed in paired samples of peripheral blood mononuclear cells (PBMC) and vitreous T cells. CD69 expression in $CD4^{+}$ T cells (n=15). Red dots indicate PBMC samples, and green dots indicate vitreous samples. Statistical significance was determined using a paired Wilcoxon signed-rank test. ***p < 0.001.

### Supplementary Figure 8

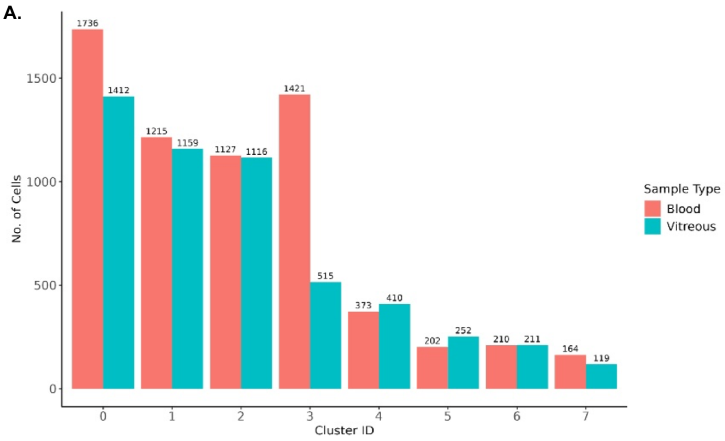


**SUPPLEMENTARY FIGURE 8: A.** Bar plot showing the distribution of CD8 T cells among different clusters (Cluster 0–6) in Blood and Vitreous samples. Each bar represents the number of cells in a given cluster. Numbers above the bars indicate the exact cell counts for each cluster.

### Supplementary Figure 9

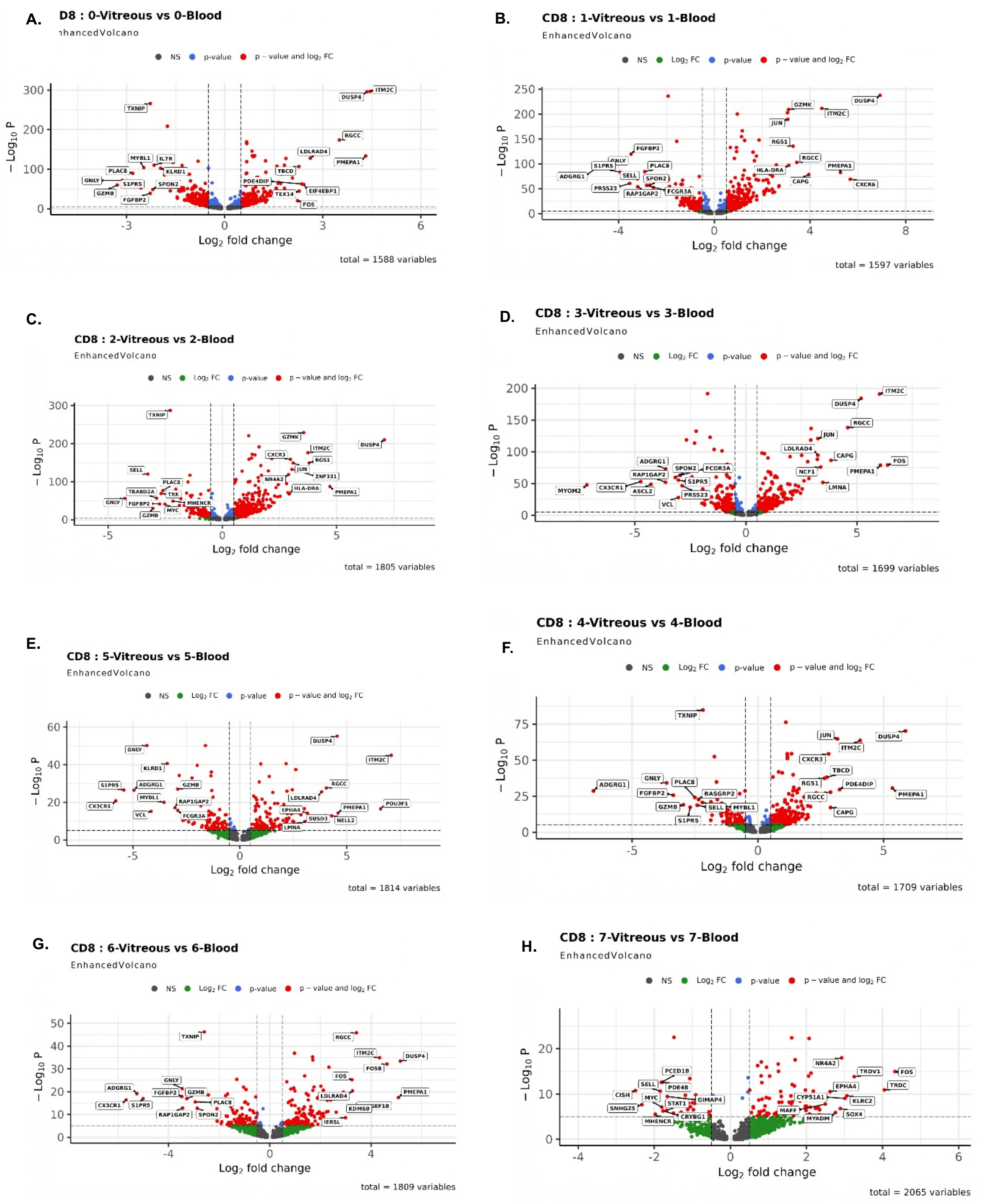


**FIGURE 9: Differential gene expression between vitreous and blood CD8⁺ T-cell subsets. A–H**. Enhanced volcano plots depicting differential gene expression between vitreous-derived and peripheral blood CD8⁺ T-cell subsets across clusters 0–7. A to H correspond to CD8⁺ clusters 0 through 7, respectively. The x-axis shows $\log_2$ fold change (vitreous vs blood), and the y-axis shows $-\log_{10}$ (p-value. Genes not meeting significance thresholds are shown in grey (NS), genes with significant fold change only are shown in green, genes with significant p-values only are shown in blue, and genes meeting both criteria (adjusted p-value and $\log_2$ fold change thresholds) are highlighted in red. Dashed vertical and horizontal lines indicate cutoff thresholds for fold change and statistical significance. The total number of genes analysed in each comparison is indicated in the lower right corner of each panel.

**Supplementary Table 2: Patient data summary for vitreous samples collected for multi-color flow cytometry evaluation**

| **Variable** | **Proportion of total cohort, n=74 (%)** |
|---|---|
| Age, median (range, years) | 38 (16 – 67) |
| Sex (males) | 35 (47.3) |
| Anatomical classification* | |
| - Intermediate uveitis | 34 (45.9) |
| - Panuveitis | 29 (31.2) |
| - Posterior uveitis | 11 (14.9) |
| Etiology | |
| - Noninfectious uveitis | 56 (75.7) |
| - Ocular tuberculosis | 18 (24.3) |
| Disease activity at sampling[#] | |
| - Moderate vitreous haze | 44 (59.5) |
| - Severe vitreous haze | 30 (40.5) |
| Systemic treatment at sampling | |
| - No treatment | 63 (85.1) |
| - Systemic corticosteroids | 2 (2.7) |
| - Systemic immunomodulatory therapy | 9 (12.2) |

*Jabs DA, Nussenblatt RB, Rosenbaum JT; Standardization of Uveitis Nomenclature (SUN) Working Group. Standardization of uveitis nomenclature for reporting clinical data. Results of the First International Workshop. Am J Ophthalmol. 2005;140(3):509-16.

[#] Nussenblatt RB, Palestine AG, Chan CC, Roberge F. Standardization of vitreal inflammatory activity in intermediate and posterior uveitis. Ophthalmology. 1985;92(4):467-71.
