## Supplemental Table for "Distinct Programs of Tissue Adaptation Shape Vitreous CD4^+^ and CD8^+^ T-cell States in Chronic Uveitis"

# CD4

| Vitreous_PBMC shared clones | | | Vitreous unique clones | | | Blood unique clones | | |
|---|---|---|---|---|---|---|---|---|
| cluster | gene | FC | Vitreous cluster | gene | FC | Blood cluster | gene | FC |
| 0-Blood | CHN1 | 5.754716981 | 0 | LST1 | 2.808239092 | 0 | CTSH | 7.077223851 |
| 0-Blood | NCR3 | 5.32722372 | 0 | CTSH | 2.457755181 | 0 | LGALS1 | 6.157184751 |
| 0-Blood | CCR6 | 4.36489854 | 0 | LGALS3 | 2.272941094 | 0 | BHLHE40 | 4.7529992 |
| 0-Blood | MHENCR | 4.281509434 | 0 | DPP4 | 2.257864425 | 0 | CCR6 | 4.226049274 |
| 0-Blood | KLF2 | 3.879275351 | 0 | HOPX | 2.212744315 | 0 | AL136456.1 | 3.878863072 |
| 0-Blood | TXNIP | 3.564767327 | 0 | FAM241A | 2.118349233 | 0 | S100A4 | 3.850460223 |
| 0-Blood | LTB | 3.422111059 | 0 | TNFSF13B | 2.09906194 | 0 | NSG1 | 3.664990923 |
| 0-Blood | RASGRP2 | 3.035455111 | 0 | CD40LG | 2.063829749 | 0 | TNFRSF4 | 3.626401916 |
| 0-Blood | IL7R | 2.655475588 | 0 | UBXN11 | 2.034771879 | 0 | TIMP1 | 3.603618375 |
| 0-Blood | TAGAP | 2.601447402 | 0 | HLA-DQA1 | 2.023888527 | 0 | RUNX2 | 3.521094045 |
| 0-Vitreous | HDGFL3 | 8.117647059 | 1 | GZMK | 4.679156783 | 1 | ICOS | 2.220779221 |
| 0-Vitreous | TRBV4-2 | 6.407960199 | 1 | RGS1 | 4.367788416 | 1 | MT2A | 2.128630705 |
| 0-Vitreous | TNFSF13B | 4.901220866 | 1 | NR4A2 | 3.120727206 | 1 | TSHZ2 | 2.116271963 |
| 0-Vitreous | CTSH | 4.630962833 | 1 | ZNF331 | 3.114523736 | 1 | FAAH2 | 2.101401365 |
| 0-Vitreous | MAL | 4.56502601 | 1 | JUNB | 3.108939409 | 1 | PELI1 | 1.938609626 |
| 0-Vitreous | NELL2 | 4.518233462 | 1 | CD27 | 2.897131251 | 1 | CEMIP2 | 1.888484848 |
| 0-Vitreous | CLU | 4.329411765 | 1 | PDCD1 | 2.745489427 | 1 | ARID5B | 1.854300856 |
| 0-Vitreous | TMIGD2 | 4.154486037 | 1 | ITM2C | 2.692556158 | 1 | RAPGEF6 | 1.845706248 |
| 0-Vitreous | MYADM | 3.794455713 | 1 | SOCS3 | 2.612270841 | 1 | SMCHD1 | 1.796997498 |
| 0-Vitreous | GLUL | 3.646637149 | 1 | ZFP36 | 2.497062809 | 1 | FYN | 1.782567646 |
| 1-Blood | IGLV2-14 | 4.987468672 | 2 | CEBPD | 1.557744541 | 2 | CD7 | 1.498019541 |
| 1-Blood | KLF2 | 2.457801564 | 2 | AOAH | 1.548966268 | 2 | CCR7 | 1.482499451 |
| 1-Vitreous | ST8SIA1 | 5.129644269 | 2 | ZEB2 | 1.434261477 | 2 | FHIT | 1.452380952 |
| 1-Vitreous | CTLA4 | 4.983870968 | 2 | CXCR6 | 1.429011479 | 2 | ATP5PO | 1.410968868 |
| 1-Vitreous | ENC1 | 4.0997543 | 2 | CCL5 | 1.405651762 | 2 | SNHG29 | 1.322180957 |
| 1-Vitreous | ICOS | 3.72594697 | 2 | ITGAE | 1.361358552 | 2 | GAS5 | 1.304212044 |
| 1-Vitreous | PDCD1 | 3.644226044 | 2 | PDE3B | 1.348337913 | 2 | C12orf57 | 1.301540201 |
| 1-Vitreous | TOX | 3.595636364 | 2 | DAPK2 | 1.334653682 | 2 | TOMM7 | 1.171556435 |
| 1-Vitreous | GPR183 | 3.382778489 | 2 | CCNH | 1.321138378 | 2 | NACA | 1.13999734 |

| | | | | | | | | |
|---|---|---|---|---|---|---|---|---|
| 1-Vitreous | DUSP4 | 3.195849802 | 2 | KLRB1 | 1.285604001 | 2 | FAU | 1.127817122 |
| 1-Vitreous | RGS1 | 3.059215442 | 3 | S100A9 | 6.343084767 | 3 | AIF1 | 2.866940389 |
| 1-Vitreous | ARID5B | 2.906628788 | 3 | IFITM3 | 3.392203796 | 3 | TACC3 | 2.680427322 |
| 2-Blood | GNLY | 15.05820025 | 3 | FTL | 1.906337505 | 3 | PCF11 | 2.174746704 |
| 2-Blood | PRSS23 | 15.03787879 | 3 | NPC2 | 1.641957576 | 3 | FHIT | 2.113585977 |
| 2-Blood | FGFBP2 | 8.874813711 | 3 | TNFSF13B | 1.502660266 | 3 | BACH2 | 1.696397589 |
| 2-Blood | SEPSECS | 8.49197861 | 3 | TUBA1B | 1.456621425 | 3 | PRMT2 | 1.613267421 |
| 2-Blood | C1orf21 | 6.161862528 | 3 | GAS5 | 1.340933372 | 3 | SARAF | 1.335601151 |
| 2-Blood | NKG7 | 5.381977671 | 3 | CTSW | 1.307995142 | 3 | EEF2 | 1.335261147 |
| 2-Blood | KLF2 | 3.070420624 | 3 | APOO | 1.264485365 | 5 | FOXP3 | 50.878125 |
| 2-Blood | TXNIP | 2.160137284 | 3 | RBFOX2 | 1.245553992 | 5 | IKZF2 | 18.23888889 |
| 2-Vitreous | TRBV2 | 3.5628778 | 4 | AL627171.2 | 2.511012815 | 5 | RTKN2 | 16.2972973 |
| 2-Vitreous | TRAV23DV6 | 3.172881356 | 5 | IL2RA | 130.0293683 | 5 | TIGIT | 12.34714286 |
| 3-Blood | CX3CR1 | 16.43724696 | 5 | FOXP3 | 62.90977011 | 5 | LAIR2 | 11.90131579 |
| 3-Blood | CST3 | 12.01183432 | 5 | RTKN2 | 59.28105122 | 5 | TAFA2 | 11.77089041 |
| 3-Blood | LIG3 | 9.759615385 | 5 | IKZF2 | 53.99199171 | 5 | CTLA4 | 8.392268041 |
| 3-Blood | GNLY | 9.534393491 | 5 | HPGD | 25.31972172 | 5 | HLA-DRB1 | 8.285496183 |
| 3-Blood | IFIT2 | 8.147157191 | 5 | LAIR2 | 22.96908442 | 5 | IL2RB | 7.665254237 |
| 3-Blood | S1PR5 | 7.807692308 | 5 | TAFA2 | 22.25345604 | 5 | STAM | 7.400454545 |
| 3-Blood | FGFBP2 | 6.662564103 | 5 | SELL | 19.34593015 | | | |
| 3-Blood | NKG7 | 6.137389973 | 5 | TIGIT | 14.02447104 | | | |
| 4-Blood | TSSK3 | 207.5 | 5 | TBC1D4 | 12.88899861 | | | |
| 4-Blood | BX323046.2 | 207.5 | 6 | FOS | 28.13803573 | | | |
| 4-Blood | C20orf144 | 207.5 | 6 | FOSB | 17.71077929 | | | |
| 4-Blood | KCTD16 | 155.625 | 6 | LMNA | 8.039582315 | | | |
| 4-Blood | AL450384.2 | 103.75 | 6 | MYADM | 6.364875486 | | | |
| 4-Blood | NAALAD2 | 103.75 | 6 | RASGEF1B | 5.170634921 | | | |
| 4-Blood | PPFIBP1 | 103.75 | 6 | DUSP1 | 4.960022135 | | | |
| 4-Blood | CEBPA | 103.75 | 6 | ARL4A | 4.61388212 | | | |
| 4-Blood | C9orf147 | 77.8125 | 6 | RGCC | 4.560783083 | | | |
| 4-Blood | AL109824.1 | 77.8125 | 6 | KLF6 | 4.558981822 | | | |
| 4-Vitreous | SLCO5A1 | 11.34598214 | 6 | PPP1R15A | 4.04066617 | | | |

| | | |
|---|---|---|
| 4-Vitreous | ZXDA | 10.74175824 |
| 4-Vitreous | PRKCA | 9.521103896 |
| 4-Vitreous | PHC1 | 6.347402597 |
| 4-Vitreous | CREB3L2 | 4.906370656 |
| 4-Vitreous | TRNAU1AP | 4.25 |
| 6-Vitreous | FADS1 | 29.35714286 |
| 6-Vitreous | MAP10 | 25.6875 |
| 6-Vitreous | AFG1L | 25.6875 |
| 6-Vitreous | TSPOAP1 | 23.35227273 |
| 6-Vitreous | AHDC1 | 22.01785714 |
| 6-Vitreous | NBR2 | 22.01785714 |
| 6-Vitreous | AL021707.1 | 22.01785714 |
| 6-Vitreous | FOSB | 20.93055556 |
| 6-Vitreous | OTUD1 | 18.68181818 |
| 6-Vitreous | FOS | 17.71551724 |

## CD8

| Vitreous_PBMC shared clones | | | Vitreous unique clones | | | Blood unique clones | | |
|---|---|---|---|---|---|---|---|---|
| cluster | gene | FC | vitreous cluster | gene | FC | blood cluster | gene | FC |
| 1-Blood | TRBV5-1 | 2.992409981 | 0 | NR4A3 | 2.069405528 | 0 | GZMK | 17.48741336 |
| 1-Blood | MYOM2 | 2.532452544 | 0 | ZNF331 | 2.054052431 | 0 | KLRB1 | 6.771872843 |
| 1-Blood | TRAV19 | 2.41114172 | 0 | JUNB | 1.975294752 | 0 | XCL2 | 6.374935377 |
| 1-Blood | TTC38 | 2.23987568 | 0 | NR4A2 | 1.797135798 | 0 | NR4A2 | 5.494606835 |
| 1-Blood | CD300A | 2.237553833 | 0 | CMC1 | 1.70103499 | 0 | RGS1 | 5.424287119 |
| 1-Blood | TRGV10 | 2.188168095 | 0 | ENC1 | 1.661992846 | 0 | ZNF331 | 4.233920547 |
| 1-Blood | CX3CR1 | 2.148717949 | 0 | IFRD1 | 1.639033575 | 0 | DUSP2 | 3.899301466 |
| 1-Blood | ASCL2 | 2.146530958 | 0 | DUSP2 | 1.611553602 | 0 | DUSP1 | 3.754043783 |
| 1-Blood | ZNF683 | 2.124507042 | 0 | MIDN | 1.548 | 0 | JUNB | 3.653692745 |
| 1-Blood | PRSS23 | 2.099447983 | 0 | SERTAD1 | 1.531424406 | 0 | CMC1 | 3.580679956 |
| 6-Blood | MYADM | 5.597496088 | 1 | GZMB | 2.371929825 | 1 | ZNF683 | 1.543852421 |
| 6-Blood | CRIP1 | 5.125084636 | 1 | AGTRAP | 1.951267057 | 1 | HLA-C | 1.30316451 |
| 6-Blood | TAGLN2 | 3.808427699 | 1 | ACAA2 | 1.921702404 | 1 | ACTB | 1.282944792 |

| | | | | | | | | |
|---|---|---|---|---|---|---|---|---|
| 6-Blood | TSPAN2 | 3.607065343 | 1 | CXCR6 | 1.870961887 | 1 | ACTG1 | 1.261123605 |
| 6-Blood | ARL4A | 3.22896625 | 1 | S100A4 | 1.848701787 | 1 | CTSW | 1.23238673 |
| 6-Blood | CCDC167 | 3.145096997 | 1 | LGALS1 | 1.787570156 | 1 | MYL12A | 1.217000938 |
| 6-Blood | LMNA | 2.984249157 | 1 | PTMS | 1.701455343 | 2 | LEF1 | 4.284311968 |
| 6-Blood | PRXL2C | 2.810101991 | 1 | IFITM2 | 1.640981108 | 2 | SERINC5 | 2.998328225 |
| 6-Blood | S100A10 | 2.624902267 | 1 | CDK2AP2 | 1.620716299 | 2 | TCF7 | 2.434989998 |
| 6-Blood | ANXA1 | 2.490888763 | 1 | ALOX5AP | 1.586289148 | 2 | CAMK4 | 2.219650747 |
| 0-Blood | KLRB1 | 3.712907738 | 2 | ZBTB20 | 1.357918745 | 2 | SELL | 2.083227597 |
| 0-Blood | TRAV12-2 | 2.982604543 | 2 | PDE3B | 1.34318555 | 2 | RUNX2 | 2.079290479 |
| 0-Blood | IL7R | 2.923289657 | 3 | GNLY | 42.30507588 | 2 | PDE3B | 1.801712363 |
| 0-Blood | TRBV2 | 2.920519022 | 3 | KLRD1 | 3.835767065 | 2 | OXNAD1 | 1.755875415 |
| 0-Blood | PIM2 | 2.526119036 | 3 | ZNF683 | 3.655432713 | 2 | IL7R | 1.698858311 |
| 0-Blood | CMC1 | 2.489689683 | 3 | HOPX | 3.536955323 | 2 | FOXP1 | 1.646762015 |
| 0-Blood | PHACTR2 | 2.482649025 | 3 | THUMPD3 | 3.421368547 | 3 | TRGV5 | 12.28951475 |
| 0-Blood | DUSP5 | 2.244394287 | 3 | GALNT2 | 2.782404854 | 3 | TRBV9 | 11.89454233 |
| 0-Blood | PLCB1 | 2.13388785 | 3 | GZMB | 2.618643297 | 3 | CSGALNACT1 | 4.431736729 |
| 0-Blood | LTB | 2.095633551 | 3 | ABCB1 | 2.49271137 | 3 | KLRC1 | 3.32765942 |
| 3-Blood | GNLY | 4.114467873 | 3 | ITGA1 | 2.41129115 | 3 | HLA-G | 3.28401142 |
| 3-Blood | TRDV1 | 3.867549943 | 3 | SLK | 2.170686456 | 3 | GNLY | 2.675512602 |
| 3-Blood | TRBV9 | 3.613392054 | 4 | AL627171.2 | 2.590623968 | 3 | TYROBP | 2.611938087 |
| 3-Blood | MYOM2 | 3.382159484 | 5 | HLA-DRA | 4.946779083 | 3 | ASCL2 | 2.328846742 |
| 3-Blood | ASCL2 | 3.087829072 | 5 | HLA-DRB1 | 3.53648524 | 3 | HOPX | 2.246886645 |
| 3-Blood | TRAV1-2 | 2.993406958 | 5 | HLA-DQA1 | 3.374544106 | 3 | FCGR3A | 2.22131886 |
| 3-Blood | PRSS23 | 2.94857749 | 5 | HLA-DQB1 | 3.090083974 | 5 | HLA-DRA | 9.702879581 |
| 3-Blood | TRGV2 | 2.937670185 | 5 | HLA-DMB | 2.937174096 | 5 | CD74 | 3.527623496 |
| 3-Blood | SYNGR1 | 2.922300442 | 5 | HLA-DRB5 | 2.842614766 | 5 | HLA-DRB1 | 3.487649965 |
| 3-Blood | GZMB | 2.77632695 | 5 | CD74 | 2.483559514 | 5 | HLA-DRB5 | 3.215800478 |
| 2-Blood | THEMIS2 | 2.557285714 | 5 | HLA-DPA1 | 2.413937151 | 5 | HLA-DPA1 | 2.161151589 |
| 2-Blood | TTC38 | 2.219733185 | 5 | HLA-DMA | 2.198093782 | 5 | HLA-DPB1 | 2.147010829 |
| 2-Blood | CEP78 | 2.214746544 | 5 | HLA-DPB1 | 1.936190561 | 6 | LMNA | 10.10573643 |
| 2-Blood | VCL | 2.126640927 | 6 | FOSB | 7.897399189 | 6 | MYADM | 9.727234519 |
| 2-Blood | ADGRG1 | 2.081046278 | 6 | FOS | 7.41402792 | 6 | CRIP1 | 6.343278264 |

| | | |
|---|---|---|
| 2-Blood | CD226 | 2.005714286 |
| 2-Blood | AGTPBP1 | 1.972699758 |
| 2-Blood | PRSS23 | 1.893320719 |
| 2-Blood | CX3CR1 | 1.83956044 |
| 2-Blood | IGF2R | 1.774865736 |
| 5-Blood | HLA-DRA | 4.349240582 |
| 5-Blood | CSGALNACT1 | 3.624740125 |
| 5-Blood | PROK2 | 2.986858418 |
| 5-Blood | HLA-DRB1 | 2.819446538 |
| 5-Blood | HLA-DQB1 | 2.78277293 |
| 5-Blood | CES1 | 2.582006664 |
| 5-Blood | TRBV9 | 2.503620975 |
| 5-Blood | HLA-DRB5 | 2.28554553 |
| 5-Blood | TRAV1-2 | 2.269554323 |
| 5-Blood | TRGV2 | 2.172059383 |
| 4-Blood | HFM1 | 4.862565653 |
| 4-Blood | AL627171.2 | 2.668345753 |
| 4-Blood | TRIM44 | 2.175622689 |
| 4-Blood | MYBL1 | 1.766451713 |
| 4-Blood | STAT1 | 1.737084576 |
| 4-Blood | TXNIP | 1.590649608 |
| 4-Blood | FLNA | 1.546875654 |
| 7-Blood | FAXDC2 | 69.28125 |
| 7-Blood | DLGAP3 | 48.19565217 |
| 7-Blood | ZFYVE9 | 44.34 |
| 7-Blood | ACVR2B | 41.05555556 |
| 7-Blood | HAPLN3 | 38.22413793 |
| 7-Blood | MAP3K9 | 34.640625 |
| 7-Blood | TMTC2 | 30.79166667 |
| 7-Blood | GSTM3 | 29.17105263 |
| 7-Blood | IGFBP3 | 27.7125 |
| 7-Blood | ADAM19 | 26.94270833 |

| | | |
|---|---|---|
| 6 | LMNA | 6.869655459 |
| 6 | MYADM | 4.730420645 |
| 6 | KLF6 | 4.383831422 |
| 6 | PPP1R15A | 3.993179023 |
| 6 | ANKRD28 | 3.599173554 |
| 6 | ARL4A | 3.332125239 |
| 6 | ATP2B1 | 3.26700227 |
| 6 | TUBA1A | 3.126272433 |
| 7 | NSMCE1-DT | 112.625 |
| 7 | ZACN | 75.08333333 |
| 7 | MAL | 73.05405405 |
| 7 | TRPC1 | 69.30769231 |
| 7 | CCNE2 | 56.3125 |
| 7 | LYPD3 | 53 |
| 7 | TRABD2A | 48.05333333 |
| 7 | NAP1L2 | 47.42105263 |
| 7 | SCML1 | 44.49382716 |
| 7 | SMPD3 | 42.9047619 |

| | | |
|---|---|---|
| 6 | SVIL | 6.060411655 |
| 6 | ANXA1 | 3.620032131 |
| 6 | TAGLN2 | 3.471007091 |
| 6 | CCDC167 | 2.924200333 |
| 6 | S100A10 | 2.873037083 |
| 6 | CA5B | 2.808535941 |
| 6 | S100A11 | 2.545508982 |
| 7 | ADAM19 | 56.73913043 |
| 7 | SGK1 | 52.72727273 |
| 7 | CARMIL1 | 51.78571429 |
| 7 | BX664615.2 | 36.25 |
| 7 | CLEC11A | 29 |
| 7 | MPP1 | 27.61904762 |
| 7 | TAF4B | 22.65625 |
| 7 | HIF1A-AS3 | 18.70967742 |
| 7 | IL12RB2 | 16.11111111 |
| 7 | TNFRSF25 | 15.42553191 |

| | | |
|---|---|---|
| 0-Vitreous | PMEPA1 | 4.984168821 |
| 0-Vitreous | DUSP4 | 4.404494705 |
| 0-Vitreous | TEX14 | 4.276682156 |
| 0-Vitreous | ITM2C | 4.063575375 |
| 0-Vitreous | ZNF331 | 3.970868578 |
| 0-Vitreous | CD27 | 3.545687274 |
| 0-Vitreous | RGS1 | 3.527635923 |
| 0-Vitreous | EIF4EBP1 | 3.399171271 |
| 0-Vitreous | ICOS | 3.231616389 |
| 0-Vitreous | PDE4DIP | 3.181177367 |
| 4-Vitreous | KIAA0513 | 4.040086207 |
| 4-Vitreous | NELL2 | 3.80474934 |
| 4-Vitreous | PDE4DIP | 3.467897092 |
| 4-Vitreous | TLE3 | 3.406299213 |
| 4-Vitreous | TRAV8-4 | 3.402678571 |
| 4-Vitreous | STRBP | 3.390282132 |
| 4-Vitreous | DUSP4 | 3.320744289 |
| 4-Vitreous | ITM2C | 3.310966415 |
| 4-Vitreous | SUSD3 | 3.055084746 |
| 4-Vitreous | EPHA4 | 2.884 |
| 3-Vitreous | RGCC | 5.129184347 |
| 3-Vitreous | LMNA | 4.633062086 |
| 3-Vitreous | CAPG | 3.965071603 |
| 3-Vitreous | TRDV1 | 3.717216117 |
| 3-Vitreous | HLA-DRA | 3.502024076 |
| 3-Vitreous | TBCD | 3.431439538 |
| 3-Vitreous | LDLRAD4 | 3.390320062 |
| 3-Vitreous | ITM2C | 3.382637098 |
| 3-Vitreous | ZNF683 | 3.237451737 |
| 3-Vitreous | DUSP4 | 3.192897498 |
| 6-Vitreous | FOS | 29.43358396 |
| 6-Vitreous | FOSB | 28.71088435 |

| | | |
|---|---|---|
| 6-Vitreous | LMNA | 14.39829424 |
| 6-Vitreous | RGCC | 11.23469388 |
| 6-Vitreous | TRAV16 | 10.01444623 |
| 6-Vitreous | AL133415.1 | 8.13546798 |
| 6-Vitreous | MYADM | 7.375882672 |
| 6-Vitreous | CFAP20 | 7.231527094 |
| 6-Vitreous | PPP1R15A | 7.044145873 |
| 6-Vitreous | HIST1H3A | 6.553571429 |
| 1-Vitreous | CAPG | 4.177654584 |
| 1-Vitreous | ITM2C | 4.13400077 |
| 1-Vitreous | PMEPA1 | 3.549141049 |
| 1-Vitreous | NCF1 | 3.385357828 |
| 1-Vitreous | DUSP4 | 3.328857143 |
| 1-Vitreous | SUSD3 | 3.248714883 |
| 1-Vitreous | CXCR3 | 3.240574601 |
| 1-Vitreous | FAM102A | 3.147008086 |
| 1-Vitreous | EIF4EBP1 | 3.116431775 |
| 1-Vitreous | PPP1R14B | 3.081399653 |
| 2-Vitreous | PMEPA1 | 3.648034854 |
| 2-Vitreous | ITM2C | 3.324090808 |
| 2-Vitreous | NELL2 | 3.309320041 |
| 2-Vitreous | SUSD3 | 3.236967987 |
| 2-Vitreous | NCF1 | 3.094509501 |
| 2-Vitreous | FAM102A | 3.007486513 |
| 2-Vitreous | LDLRAD4 | 2.996387899 |
| 2-Vitreous | MLLT3 | 2.95702533 |
| 2-Vitreous | DUSP4 | 2.871746295 |
| 2-Vitreous | DUSP16 | 2.847232711 |
| 5-Vitreous | HLA-DRA | 8.475632044 |
| 5-Vitreous | HLA-DMB | 7.575 |
| 5-Vitreous | SUSD3 | 4.695238095 |
| 5-Vitreous | ABCG1 | 4.551942618 |

| | | |
|---|---|---|
| 5-Vitreous | HLA-DMA | 4.370846224 |
| 5-Vitreous | PMEPA1 | 4.265533981 |
| 5-Vitreous | HLA-DRB1 | 4.251638402 |
| 5-Vitreous | DUSP4 | 3.962503786 |
| 5-Vitreous | DUSP10 | 3.938151261 |
| 5-Vitreous | CD74 | 3.721593579 |
